# SM-Omics: An automated platform for high-throughput spatial multi-omics

**DOI:** 10.1101/2020.10.14.338418

**Authors:** Sanja Vickovic, Britta Lötstedt, Johanna Klughammer, Åsa Segerstolpe, Orit Rozenblatt-Rosen, Aviv Regev

## Abstract

The spatial organization of cells and molecules plays a key role in tissue function in homeostasis and disease. Spatial Transcriptomics (ST) has recently emerged as a key technique to capture and positionally barcode RNAs directly in tissues. Here, we advance the application of ST at scale, by presenting Spatial Multiomics (SM-Omics) as a fully automated high-throughput platform for combined and spatially resolved transcriptomics and antibody-based proteomics.

## Introduction

The spatial organization of cells and molecules is fundamental to physiological function and disease pathology, and imaging the position and level of molecules is a cornerstone of both basic biology and clinical pathology. Because gene expression is regulated at multiple levels from transcription to protein degradation, protein and RNA levels convey distinct information on gene function and cell state, as has been shown in diverse contexts including dynamic responses[1,2], in genetic variation[3], in human malignancies[4], and in single cells in suspension[5]. Single cell genomics and multi-omics approaches, such as single cell and single nucleus RNA-Seq[6–11] and CITE-Seq[5,12], have been tremendously successful at profiling diverse molecular profiles at the level of individual cells and nuclei, but typically do not preserve spatial information. The importance of studying cells in their native environment has been shown in many processes, from normal organ development to spatial deregulation in diseases and often highlighted in the context of cancer propagation and resistance to therapy[13,14].

Recent progress in spatial *in situ* profiling methods has opened the way for comprehensive profiling of location and expression simultaneously[15–27]. For spatial RNA measurements, Spatial Transcriptomics (ST)[24,26] has emerged as a versatile approach for spatial RNA profiling. In ST, a fresh-frozen tissue section is placed on top of barcoded DNA primers attached to a glass surface[24]. Following tissue staining and histological imaging, cells are permeabilized, mRNAs are spatially tagged directly in tissues and a cDNA sequencing library is generated. After sequencing, the RNA-Seq information is traced back to the spatially barcoded positions on the glass slide providing a global spatial tissue profile. ST has been applied to diverse systems and tissue types, such as brain, heart, spinal cord, melanomas, breast cancer and prostate cancer[24,28–35]. However, barriers around throughput, resolution, and efficiency[36], limit its application at large scale. In parallel, there have been advances in multiplex protein measurements *in situ* based on reading out multiple fluorescent-, heavy metal- or barcode coupled antibody tags at a time[19,20,37–40]. Some methods rely on cyclic immunostaining or *in situ* sequencing barcoding schemes, whereas others use expensive machinery for Multiplexed Ion Beam Imaging or Imaging Mass Cytometry. Few methods have combined RNA and antibody-based measurements[41,42]

To bridge this gap and make molecular tissue profiling a widely available and robust tool, we have developed Spatial Multi-Omics (SM-Omics), an end-to-end framework that uses a liquid handling platform for high-throughput combined transcriptome and antibody-based spatial tissue profiling with minimum user input and available laboratory instrumentation[43,44]. SM-Omics allows processing of up to 96 sequencing-ready libraries, of high complexity, in a ~2 days, making it the first truly high-throughput platform for spatial multi-omics.

## Results and Discussion

We devised SM-Omics for high throughput combined transcriptomics and antibody-based measurements. SM-Omics can be used for either Spatial Transcriptomics alone, or, in combination with fluorescently or DNA-barcoded antibodies to simultaneously measure spatial profiles of RNAs and proteins. Briefly, in SM-Omics, after tissue staining for traditional histology (H&E), immunofluorescence or using DNA-barcoded antibodies, glass slides are loaded into the SM-Omics platform, where, using a liquid handler robot, cells are permeabilized, mRNAs and/or antibody barcodes are spatially tagged and converted into a sequencing-ready library (**Fig1a**). The process consists of three main parts with designed stopping points to either store the processed material or load required reagents for the upcoming reactions. The first step consists of all *in situ* enzymatic reactions on the SM-Omics slide, including tissue permeabilization after staining and reverse transcription with simultaneous release of spatial capture probes (**Fig1a, I**). Each such *in situ* run holds up to 4 slides with tissues, with the number of active areas with spatial probes per slide ranging from one to 16 per slide. The second and third steps consist of RNA-Seq library preparation in standard 96 well plates, where the user can choose to run between 1 and 96 samples in parallel in 8-step increments with adjusted library consumable usage to alleviate costs. The input to these is *in situ* tissue cDNA or antibody tag material collected from SM-Omics slides in the first step, which are then processed to amplify cDNA using a T7 *in vitro* transcription approach (for cDNA) or standard PCR amplification (for antibody tags), followed by a final conversion of the amplified RNAs into sequencing-ready libraries (**Fig1a, II-III)**.

**Fig1.**
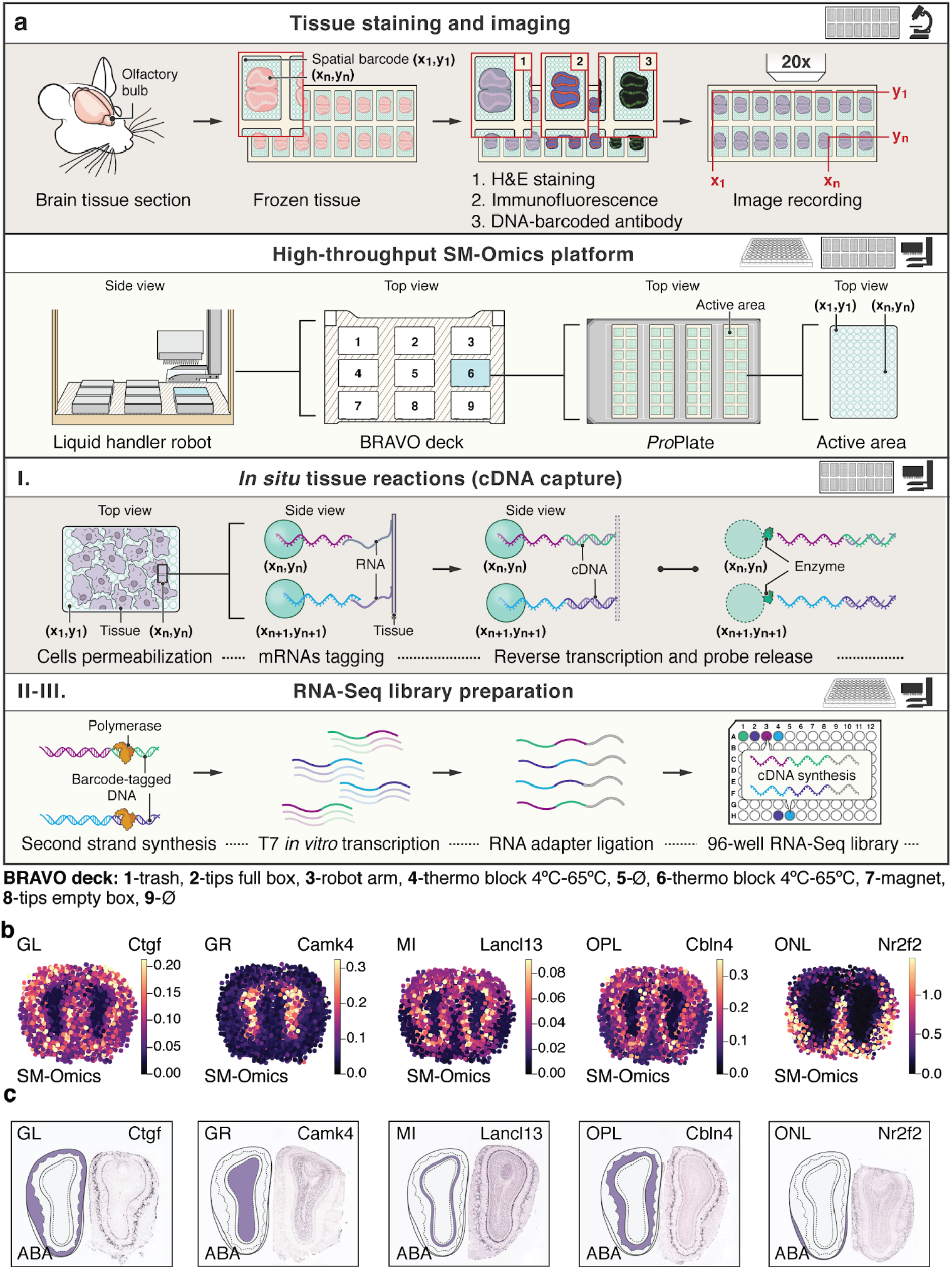
SM-Omics method creates tissue specific spatial gene expression patterns. **(a)** SM-Omics approach combines automated imaging of H&E, IF stained or tissue sections stained with DNA-barcoded antibodies with high-throughput liquid handling to create spatially resolved RNA-seq and/or antibody-seq libraries. The RNA-seq protocol consists of three main steps. (I) *in situ* reactions on a ST slide that include tissue permeabilization, capture of mRNAs on the spatial array followed by a reverse transcription reaction in solution. The transcribed material is then collected and a two-step library preparation protocol (II-III) is run in standard 96-well plates. **(b)** Examples of SM-Omics spatial gene expression patterns (color scale) detected in each of the major histological regions in the main olfactory bulb of an adult mouse brain and **(c)** corresponding ISH images from ABA for the same genes as in **(b)** with illustrated and highlighted region annotation patterns. Annotated region abbreviations: GL (Glomerular Layer), GR (Granular Cell Layer), MI (Mitral Layer), OPL (Outer Plexiform Layer) and ONL (Olfactory Nerve Layer) are shared between the panels.

SM-Omics introduces four key enhancements compared to ST: (**1**) Automation, requiring minimal user intervention; (**2**) throughput, allowing processing of 96 samples in a 2-day cycle; (**3**) enhanced quality, reflected by higher complexity RNA-Seq libraries and (**4**) combining RNA-Seq measurements with proteomics measurements including immunofluorescent (IF) staining and antibody-barcoding strategies. We first describe the core approach in the context of spatial RNA measurements (**Fig1a, II-III)**, and then its extension to include spatial protein measurements.

To test the performance of SM-Omics for spatial transcriptomics, we assessed the feasibility, reproducibility and efficiency of RNA data in two key steps, testing on the mouse olfactory bulb (MOB) and mouse cortex: (1) *in situ* tissue reactions (cDNA capture) and (2) library (RNA-Seq) preparation.

SM-Omics had enhanced performance in terms of *in situ* reactions compared to standard ST, with minimal lateral diffusion and comparable and reproducible cDNA signal intensity. Specifically, we first ran *in situ* reactions on the glass surface in optimization mode, where cDNA molecules are *in situ* fluorescently labeled to create a spatial cDNA footprint[35] (**FigS1a**). We compared the localized cDNA footprint to the histological H&E pattern and measured the lateral tissue permeabilization effects. This provides an optimal set of parameters needed to successfully run tissue-specific reactions and to ensure minimal lateral cross-talk between adjacent spatial measurements. Testing on the adult mouse cortex (**FigS1b-e**) showed that SM-Omics resulted in no mixing of material between spatial measurements with no lateral diffusion (mean −0.06 μm ±0.51 sd), which is 4X weaker lateral diffusion signal than in ST performed on adjacent tissue sections (p<0.01, two-sided *t*-test, **FigS1f,g**), and 30x weaker diffusion signal compared to previous reports[24,35,45]. Moreover, the signal intensity of the fluorescent cDNA footprint was highly reproducible within and between SM-Omics runs: there were no significant differences (Wilcoxon rank-sum test, p>0.05) between the cDNA signal intensities from adjacent adult mouse main olfactory bulb (MOB) tissue replicates on a single glass slide (n=3), single run (n=3) or separate runs (n=3) (**FigS2**). SM-Omics yielded robust spatial fluorescent patterns in three other tissues: mouse cortex, a mouse model of colorectal cancer, and a distal part of the mouse colon (**FigS3**).

To process the generated data efficiently, we also developed SpoTteR, a fast and fully automated end-to-end image integration method. With SpoTteR, images are automatically downscaled and barcode spots positions reconstructed using iterative blob detection and grid fitting (**Methods**), accounting for common imaging artifacts, such as uneven tissue coloration or pipetting bubbles. SpoTteR then registers tissue coordinates through a masking process to produce a gene-by-barcode matrix overlaid on top of morphological features (**FigS4**). Compared to manual and semi-automated approaches[46] SpoTter is up to 14X faster with low false discovery rates (FP 3.54% and FN 1.18%, *vs*. >15% of grid spots as FNs in other approaches[46]; **FigS5**), when applied to human lung cancer, human arthritis and mouse colon data.

Using the SM-Omics end-to-end toolbox (**Fig1a**) we prepared and sequenced 18 SM-Omics libraries from the main olfactory bulb of the adult mouse brain, and compared them to standard ST libraries. SM-Omics libraries were more sensitive than ST, with a 58% higher number of protein-coding genes (4,369 genes), and 1.5-fold higher number of unique transcripts (UMIs) (Wilcoxon’s rank-sum test, `p≤0.05`, **Methods, FigS6a,b**). Both ST and SM-Omics had similar correlations between their respective pseudo-bulk averages and replicates (**FigS6c**), but SM-Omics exhibited an increase on average (n=3) in the number of transcripts captured in more than half of the annotated morphological regions (**Methods**, **FigS6d**).SM-Omics (n=3) also performed comparably to newer generation array designs (n=3) (Visium, 10X Genomics) in detected genes and UMIs per measurement (p≥0.05, Wilcoxon’s rank sum test) in the adult mouse brain cortex tissues (**FigS6e-g**). We also confirmed that our liquid handling system processed standard spatial library preparations robustly with no significant variation (Wilcoxon’s rank-sum test, p≥0.05) between runs (**FigS7a-b**). This increased efficiency in SM-Omics, as reflected in the number of genes and UMIs detected per (x,y) coordinate, was due to several optimizations in library preparations. First, we introduced simultaneous release of barcoded primers and capture of mRNA molecules (**Methods**). This hybrid can then be used as a template in the reverse transcription reaction in solution instead of solid surface as previously performed; this also decreased total processing time from ~1.5 days to ~6h. Second, we improved the efficiency of library preparation `reactions, by increasing the amount of sequencing adaptors and reaction time for adaptor ligation to the template (Wilcoxon’s rank sum test, p≤0.05)` (**FigS7c-d**).

We also compared SM-Omics and ST in terms of specific detection of known and novel specific spatial expression patterns. We used Splotch[31,47] to align our replicate tissue sections and generate posterior spatial gene expression estimates. We confirmed that region-enriched and upregulated genes were present in the major spatial layers (**Methods**) of the MOB compared to the Allen Brain Atlas[48] (**FigS8a,b**). While known gene patterns detected as layer-enriched agreed between SM-Omics and ST (**FigS8c-f**), SM-Omics’s overall specificity was higher (**FigS8a**). The increased sensitivity at the same sequencing depth (by down-sampling, **Methods**), allowed us to reproducibly measure the spatial gene expression of newly detected targets, such as Ctgf in the Glomerular Layer, Camk4 in the Granular Cell Layer, Lancl3 in the Mitral Layer and Cbln4 in the Outer Plexiform Layer (**Fig1b,c**).

Next, we implemented a combined spatial transcriptomics and antibody-based read-out into our fully automated spatial multi-omics platform, by using either immunofluorescence and imaging or DNA-barcoding and sequencing.

We first developed a protocol that combined antibody-based immunofluorescence (IF) with spatial transcriptomics (**Fig2a**, **Methods**). Localized cDNA footprints after nuclear (DAPI) and IF stainings of the tissue (**Fig2b**, **FigS9a**) showed that mRNAs were laterally diffusing only 0.16±1.21μm outside of the nucleus, again indicating minimal lateral cross-talk between adjacent spatial measurements. We next created SM-Omics mouse brain cortex libraries following immunostaining with an antibody against the brain protein NeuN, which is highly expressed in most neuron nuclei (**Fig2c**). Library complexities, signal specificity and RNA expression patterns were similar to those in standard (H&E stained) ST measurements and in the Allen Brain Atlas[48] (**FigS9b-d**), confirming that our protocol for simultaneous immunofluorescent and transcriptome measurements provided high-quality mRNA data. Next, comparing the antibody IF signals and corresponding RNA expression (**Fig2c**), there was significant correlation between NeuN mRNA and protein expression (Spearman’s ρ 0.73, `p-value≤0.05`, **Fig2c**). Notably, in some regions (*e.g.*, hypothalamus) RNA expression was low but protein expression was substantial (**Fig2c**). This may be due to either a biological difference, or to the differences in sensitivity and saturation of RNA-Seq *vs*. IF.

**Fig2.**
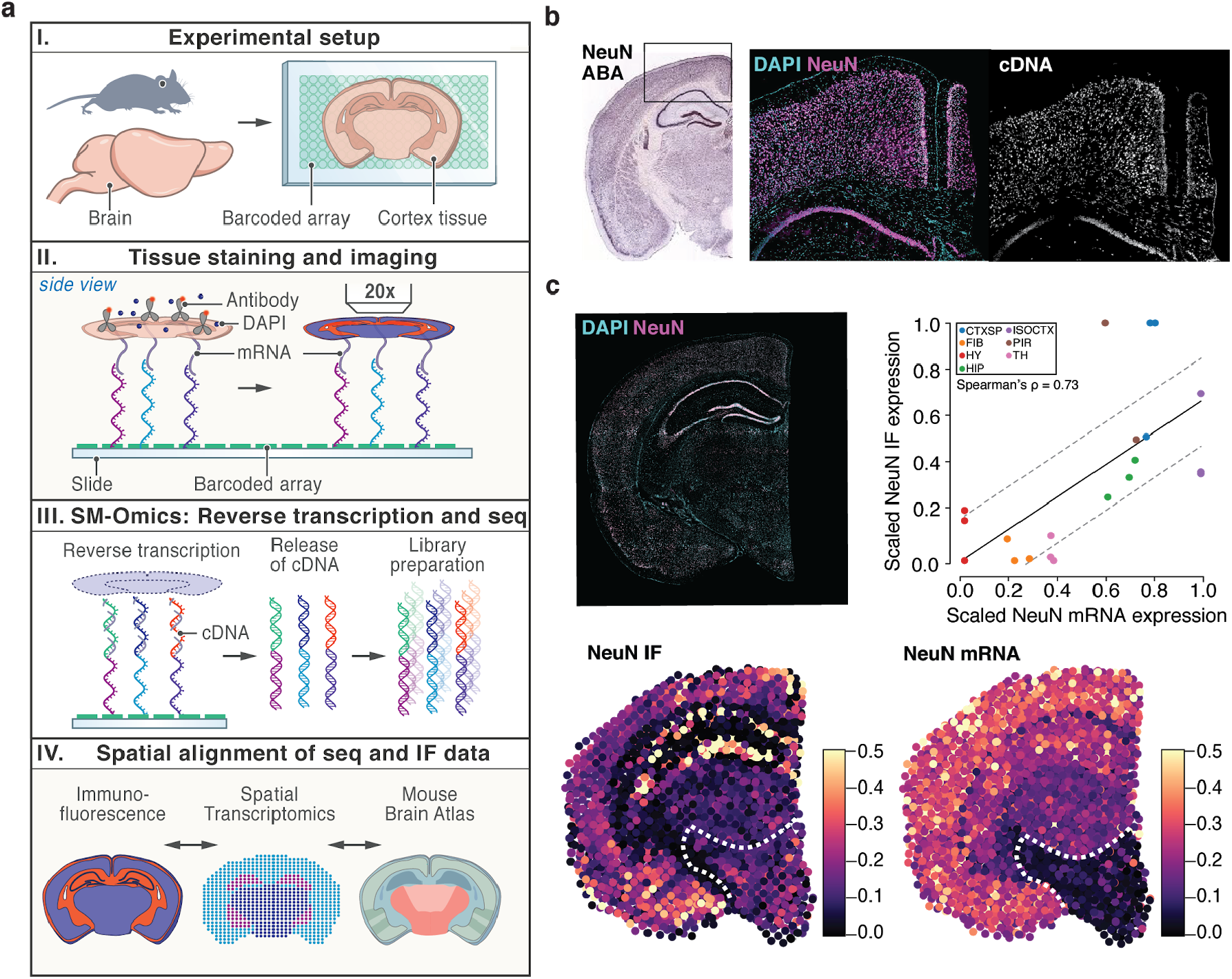
SM-Omics method tissue specific gene and IF antibody expression patterns. **(a)**Tissue sections are placed on the spatial array (I), stained for nuclear and corresponding antigen targets, imaged for fluorescent IF signals (II) and SM-Omics libraries created (III). Spatial gene and antibody expression data are processed and compared to the reference ABA atlas (IV). **(b)**Neuronal target NeuN; was stained for antibody IF and DAPI and corresponding gene activity labeled (cDNA) confirming feasibility of *in situ* reaction conditions for IF staining. ABA reference image for the same target with labeled zoomed-in area (isocortex). **(c)**Fluorescently stained tissue section (upper left) could be analyzed and protein IF signals (color scale) aggregated in SM-Omics-like spots (NeuN IF; lower left) for comparison to mRNA expression signals (NeuN mRNA; lower right). White dashed lines circle lower part of the mouse brain *ie*. hypothalamus. Correlation between scaled NeuN IF and respective NeuN mRNA expression per tissue section (n=3; upper right). Each dot represents the mean scaled signal of all SM-Omics spatial measurements in that annotated region (color code). Line (black) represents the linear regression line between the conditions with respective standard deviations (gray dashed lines). Annotated region abbreviations: CTXSP (Cortical subplate), FIB (Fiber tracts), HY (Hypothalamus), HIP (Hippocampal formation), ISOCTX (Isocortex), PIR (Piriform areas) and TH (Thalamus).

Finally, we introduced an antibody DNA-barcoding system[5] compatible with spatial transcriptomics to increase multiplexing capacities otherwise limited with spectral overlap in imaging approaches (**Fig3a**). We tag each of 6 antibodies[5] with an amplification primer and an individual barcode tag followed by a poly(d)A sequence for capture on a poly(d)T spatially barcoded array (**Methods**). We used a similar tissue staining protocol as that for immunofluorescence, where the tissue was first *in situ* fixed with paraformaldehyde to ensure specific antigen coupling, followed by antibody staining, tissue permeabilization and SM-Omics library preparation (**Fig3a**). To benchmark our approach, we incubated adult mouse spleen tissue sections with both a fluorescently labeled antibody and a barcoded antibody, allowing us to simultaneously validate and directly compare both detection methods. We imaged the fluorescently labeled epitopes prior to any *in situ* enzymatic reactions on the array surface, coupled the antibody tags to the spatial array, such that they were copied into a stable covalent complex, while mRNA was spatially captured and transcribed on the array (**Fig3a**). We first tested a two-antibody cocktail targeting F4/80 (staining splenic red pulp macrophages) and IgD (staining marginal zone B cells in the white pulp) (**Fig3b**). We obtained high quality antibody tag (mean±sd 142±15 UMIs per SM-Omics measurement; n=3) and cDNA libraries (1,375±181 UMIs per SM-Omics measurement, n=3), with highly specific antibody tag patterns (**Fig3b**) that were well-correlated to the corresponding IF intensities across all major splenic regions (**FigS10a**, on average 78%). RNA and antibody tag levels were in agreement for IgD (Spearman’s ρ = 0.74, `p-value≤0.05` across all spatial measurements), and less so for F4/80 (Spearman’s ρ = 0.65, `p-value≤0.05` across all spatial measurements) (**FigS10b**). Finally, an SM-Omics experiment with 6 validated[49] barcoded antibodies targeting F4/80, IgD, CD163, CD38, CD4, and CD8a (**FigS10c**) successfully combined spatial transcriptomics and protein estimates in a highly multiplexed manner (**Fig3c**). CD4 and CD8 proteins (by antibody signal) and their corresponding mRNAs were spatially localized in the PALS zone, whereas IgD and CD38 protein and mRNA were enriched in the B follicles. F4/80 protein and mRNA were localized to the red pulp, but the corresponding mRNA (Adgre1) was also enriched in the marginal zone. Finally, CD163 was differentially expressed, as expected, in the red pulp with Cd163 mRNA also high in PALS.

**Fig3.**
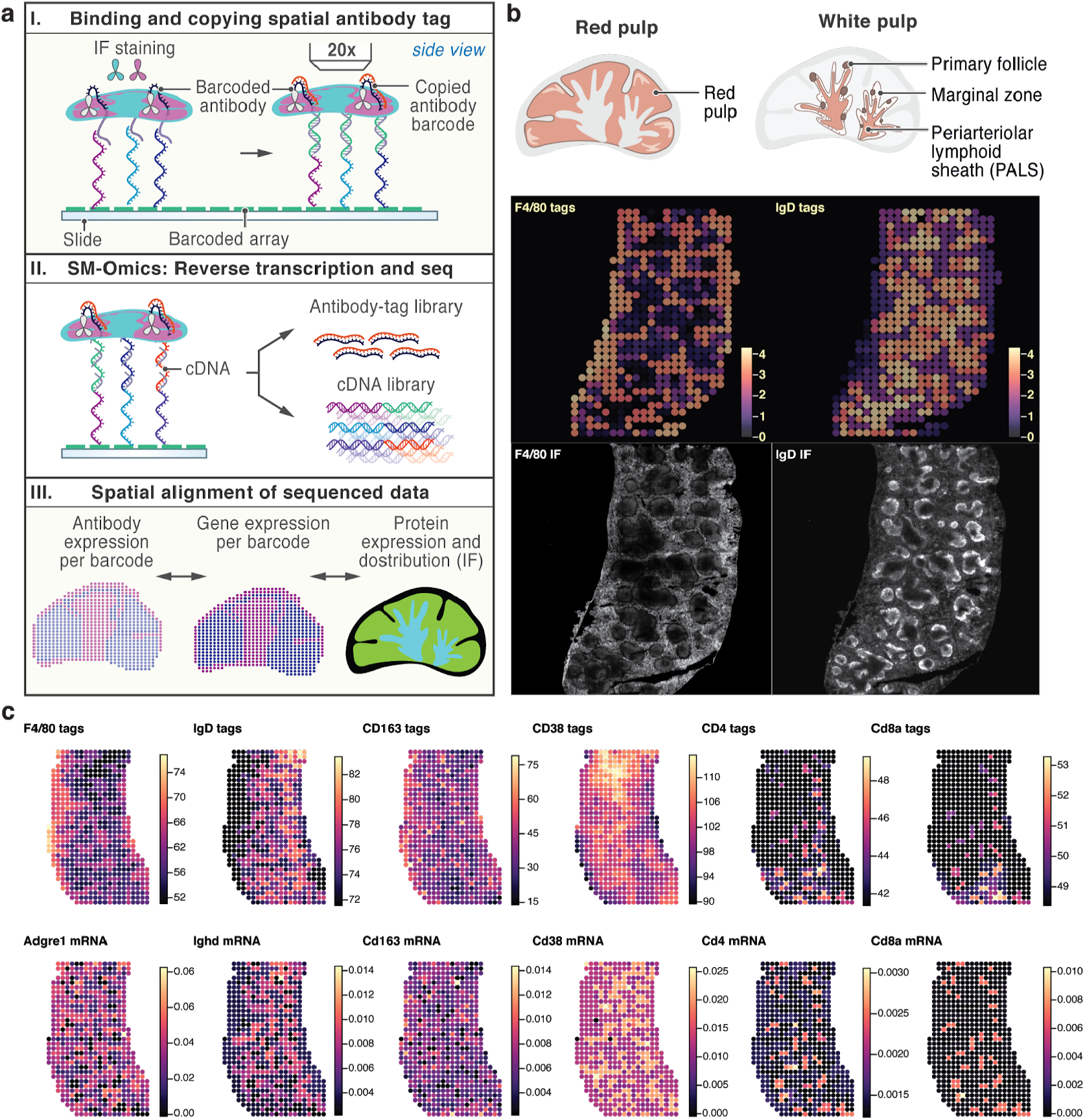
Highly-multiplexed SM-Omics tissue specific gene and antibody expression patterns. **(a)** SM-Omics approach combines automated imaging of IF antibody stained tissue sections, tagging antigens spatially *in situ* using barcoded antibodies and capturing mRNA on a spatially barcoded poly(d)T array. Frozen tissue sections are placed on a SM-Omics array, tissues stained with both IF and DNA-tagged antibodies, imaged and *in situ* copying reactions performed and at the same time as cDNA made (I). Then, both the antibody tags and cDNAs are used in the library preparation reactions and sequenced (II). Finally, spatial IF, antibody tag and gene expression patterns can be evaluated (III). **(b)** Splenic tissue illustration of red and white pulp structures followed by spatial expression profiles of sequenced antibody tags as well as IF images in splenic tissue for F4/80 staining red pulp macrophages and IgD staining marginal zone B cells in the white pulp. **(c)** Spatial expression profiles (color scale) for a 6-plex SM-Omics reaction with F4/80, IgD, CD163, CD38, CD4 and CD8a DNA-barcoded antibody-based expression in the top panel and respective gene expression shown in the bottom panel.

## Conclusions

SM-Omics is an efficient and automated workflow for combined and spatially resolved transcriptomics and antibody-based proteomics, adaptable to new array versions and designs. SM-Omics provides a more detailed molecular high-plex multi-omics characterization of tissues *in situ* and is the first high-throughput automated system for quantifying the spatial transcriptome and antibody-based proteome, by either immunofluorescence or using DNA barcoded antibodies. We confirmed SM-Omics as a robust system that can reconstruct specific cell associations across morphological layers[50,51], and characterize tissue niches in combination with antibody staining, which provide higher resolution views independently of or in combination with spatial transcriptomics patterns. Its automation on a widely-used platform enables use of appropriate study design while minimizing technical variation, and allowing broad adoption. SM-Omics does not rely on any customized microfabrication, uses commercially, widely-available liquid handlers and reagents with minimum preparation time per run (~30 min), has an end-to-end image-integrated data analysis pipeline and is readily deployable to the wide scientific community.

## Materials and methods

### Bravo system requirements

Bravo Automated Liquid Handling Platform (Agilent Technologies, USA) was equipped with a 96LT pipetting head (G5498B#042, Agilent Technologies, USA) and two Peltier thermal stations (CPAC Ultraflat HT 2-TEC, #7000166A, Agilent Technologies, USA) with PCR adapter having a mounting frame at positions 4 and 6 on the Bravo Deck and connected to an Inheco MTC Controller. On position 7, we recommend the MAGNUM FLX™ Enhanced Universal Magnet Plate (#A000400, Alpaqua, USA) to serve for magnetic bead-based clean ups. In addition, a BenchCel NGS Workstation (Front-load rack at 660 mm height) and BenchCel Configuration Labware MiniHub (option #010, Agilent Technologies, USA) were included in the automation platform setup. In case *in situ* reactions were performed, the PCR adapter was removed from position 6 to be replaced with Aluminum Heat Transfer Plate (#741I6-GS-4, V&P Scientific, Inc, USA).

### Sample collection and cryosectioning

A small piece of freshly collected tissue (~25-50 mg, about 5×5 mm) was placed on a dry and sterile Petri dish, which was placed on top of wet ice. The tissue was then very gently moved using forceps and placed on another dry part of the Petri dish to ensure little liquid was present around the tissue. The bottom of a cryomold (5×5mm, 10×10mm or 25×20mm) was filled with pre-chilled (4°C) OCT (Tissue-Tek; Sakura Finetek, USA) and the tissue transferred with forceps into the OCT-prefilled mold. The entire tissue surface was covered with pre-chilled OCT. The mold was then placed on top of dry ice and allowed the tissue to freeze for up to 5 minutes until OCT has turned completely white and hard. The tissue cryomolds were stored at −80°C until use. For cryosectioning, the ST slide and the tissue molds first reached the temperature of the cryo chamber. The OCT-embedded tissue block was attached onto a chuck with pre-chilled OCT and allowed to freeze ~5-10 min. The chuck was placed in the specimen holder and adjusted the position to enable perpendicular sectioning at 10μm thickness. Sections were gently transferred to a ST array[24] and then the back side of the slide was warmed ~10-15 sec with a finger. ST slides with tissue sections on top could be stored at −80°C for up to 6 days.

### Tissue fixation and H&E staining

The ST slide with the tissue section was warmed to 37°C for 1 minute on a thermal incubator (Eppendorf Thermomixer Option C, Germany). The tissue was then covered with 4% formaldehyde (Sigma-Aldrich, USA) in 1X PBS (Thermo Fisher Scientific, USA) for 10 minutes at room temperature (RT). The whole slide was then washed in 1X PBS in a vertical orientation to be placed back on a horizontal place for drying. 500μl isopropanol covered the tissue and ensured drying. The slide was put into an EasyDip Slide Jar Staining System (Weber Scientific) holder and the same system used for H&E staining. Five ~80 ml containers were prepared with Dako Mayers hematoxylin (Agilent, USA), Dako Bluing buffer (Agilent, USA), 5% Eosin Y (Sigma-Aldrich, USA) in 0.45M Tris acetate (Sigma-Aldrich, USA) buffer at pH 6 and two jars with nuclease-free water (ThermoFisher Scientific, USA). The slide rack was fully immersed in hematoxylin for 6 minutes and then washed by dipping the slide rack in a nuclease-free water jar 5 times following another destaining wash by dipping the slide rack in 800mL nuclease-free water for 30 times. The slide rack was put into the Dako bluing buffer and incubated for 1 minute. The slide was again washed by dipping the rack 5 times in the second nuclease-free water jar. The slide rack was finally put into the eosin and incubated for 1 minute to be washed by dipping the rack 7 times in the second water jar. The slide was removed from the rack to allow it to dry.

### Tissue fixation and IF staining

The ST slide with the tissue section was warmed to 37°C for 4 minutes on a thermal incubator (Eppendorf Thermomixer Option C, Germany) and *in situ* fixed and washed as described above. The slide was then mounted in the plastic slide holder (ProPlate Multi-Array slide system; GraceBioLabs, USA) compatible with the Aluminum Heat Transfer Plate (#741I6-GS-4, V&P Scientific, Inc, USA) on position 6 on the Bravo deck. All following antibody incubations were performed at 4°C. First, the tissues were blocked with the TruStain FcX™ PLUS (anti-mouse CD16/32, Biolegend, USA) antibody (1:100 dilution) in 0.5% Triton X-100 (Sigma-Aldrich, USA) for mouse brain tissues and 1% saponin (Sigma-Aldrich, USA) supplemented with 5% FBS (ThermoFisher Scientific, USA) for splenic tissues. This simultaneous blocking and permeabilization step lasted for 30 min. Next, the slide was washed 3x with 1x PBS (ThermoFisher Scientific, USA). After discarding the last wash, the slides were incubated with 1x PBS for 2 min. Then, antibodies were added at 1:100 dilution in 1% saponin (Sigma-Aldrich, USA) supplemented with 5% FBS (ThermoFisher Scientific, USA) for 90 min. The complete list of antibody clones and suppliers is available in **Supplementary Table 1**. The slide was again washed in the same fashion and counterstained with DAPI (Sigma-Aldrich, USA) diluted 1:1000 in 0.5% Triton X-100 (Sigma-Aldrich, USA) for 5 min. In case the reactions were performed on a SM-Omics array and not a mock polyd(T) array, the DAPI reaction was also supplemented with a Cy3 labeled anti-frame DNA probe (5’-Cy3-GGTACAGAAGCGCGATAGCAG-3’, IDT, USA) at 10 nM concentration. In case DAPI counterstaining was not used, the step was skipped. This was followed by another wash cycle. The slides were then air dried and mounted with 85% glycerol prior to imaging.

### Tissue fixation and DAPI-only staining

Similarly to performing *Tissue fixation and IF staining*, tissue sections were attached to slides and *in situ* fixed. The slide was then mounted in the plastic slide holder (ProPlate Multi-Array slide system; GraceBioLabs, USA) and all reactions performed at 4°C. Tissues were first incubated with 0.5% Triton X-100 (Sigma-Aldrich, USA) for 25 min. Next, the slide was washed 1x PBS (ThermoFisher Scientific, USA) and the tissue stained with DAPI (Sigma-Aldrich, USA) diluted 1:1000 in 0.5% Triton X-100 (Sigma-Aldrich, USA) for 15 min. If the reactions were performed on a SM-Omics array and not a mock polyd(T) array, the DAPI reaction was also supplemented with a Cy3 labeled anti-frame DNA probe (5’-Cy3-GGTACAGAAGCGCGATAGCAG-3’, IDT, USA) at 10 nM concentration in order to facilitate image registration to the SM-Omics array coordinates. This was followed by another wash cycle. The slides were then air dried and mounted with 85% glycerol prior to imaging.

### Automated imaging

Images of stained H&E tissue sections on the ST slides were taken using a Metafer Vslide scanning system (MetaSystems, Germany) installed on an Axio Imager Z2 microscope (Carl Zeiss, Germany) using an LED transmitted light source and a CCD camera (BF scanning). All images were taken with the A-P 10x/0.25 Ph1 objective lens (Carl Zeiss, Germany). For fluorescent scanning, a PhotoFLuor LM-75 lightsource (89North, USA) was used in combination with a Plan-APOCHROMAT 20x/0.8 objective (Carl Zeiss, Germany). A configuration program was made to enable automatic tissue detection, focusing and scanning on all ST arrays present on a glass slide. In short, tissue detection was based on contrast as compared to normalized background in RGB channels. Upon finding maximum contrast in a 12-step spiral-like search window field of view (FOV) pattern, the automated focal alignment in every second of each FOV (4096×3000 px) was initiated. The alignment search considered the maximum contrast z-position as in-focus using 5μm stage intervals (n=19 focal planes). The BF scanning of the predefined ST array areas was done in a total of 48 FOVs and ~30sec in 3 channels (RGB); or fluorescent scanning of 228 FOVs and ~6min for 3 fluorescent channels. Images were stitched using 60μm overlap and linear blending between FOVs with the VSlide software (v1.0.0) and then extracted using jpg compression. Multiple ST slides can be processed in the same manner without any user input for a total of 6min processing time per H&E stained slide (3 channels) or 45min for fluorescently stained slide (3 channels), including image stitching.

### SM-Omics automation

The SM-Omics protocol is divided into three main parts. The first part (1) processes all *in situ* reactions on a ST slide: tissue pre-permeabilization, permeabilization, reverse transcription with or without the release of the spatial capture probes and tissue removal. This material is collected to a standard 96-well PCR microplate (Eppendorf, Germany) and all of the following reactions (protocols 2 and 3) are run in 96-well plates. The second protocol (2) contains second strand synthesis reaction, cDNA bead purifications and T7 *in vitro* transcription. The third protocol (3) includes aRNA adapter ligation, bead purifications and second cDNA synthesis. The material is then quantified using a standard qPCR protocol and the libraries accordingly indexed for Illumina sequencing.

### Reference material preparation

In order to test reproducibility of library preparation reactions, we prepared reference material as input. 7.5μg of universal mouse reference RNA (#740100, Agilent Technologies, USA) was fragmented using NEBNext Magnesium RNA fragmentation module (NEB, USA) for 1 minute at 94°C. The sample was purified with a MinElute Cleanup kit (Qiagen, Germany) according to manufacturer’s instructions and RNA concentration and size were assessed using a Qubit RNA HS kit (ThermoFisher Scientific, USA) and Bioanalyzer Pico 6000 kit (Agilent Technologies, USA), respectively. ~2μg of fragmented RNA was incubated with either 3.3μM custom hexamer primer GACTCGTAATACGACTCACTATAGGGACACGACGCTCTTCCGATCTNNNNNNNN, T7handle_IlluminaAhandle_hexamer) or poly(d)T primer (T7handle_IlluminaAhandle_hexamer_20TVN) in the presence of 0.8mM dNTP (ThermoFisher Scientific, USA) at 65°C for 5 minutes. First strand reverse transcription was performed with a final concentration of 1X First Strand Buffer, 5 mM DTT, 2U/μl RNaseOUT and 20U/μl of Superscript III (all from Thermo Fisher Scientific, USA). The reaction was incubated at 25°C for 10 min (when using hexamer priming), followed by 50°C for 1 hr and 70°C for 15 minutes or 50°C for 1 hr and 70°C for 15 minutes for poly(d)T priming. The reaction was purified with AMPure XP beads (Beckman Coulter, USA) at a beads/DNA ratio of 0.8:1. The concentration of the material was measured on a Qubit RNA HS kit (ThermoFisher Scientific, USA) and diluted in EB (Qiagen, Germany). A release mixture of ~100ng (hexamer priming) or ~200ng (poly(d)T priming) first strand cDNA, 1X Second strand buffer (ThermoFisher Scientific, USA), 0.2μg/μl BSA and 0.5mM dNTP (ThermoFisher Scientific, USA) was used to test all library preparation reactions. Hexamer primed cDNA was used to test the reproducibility and poly(d)T primed cDNA was used to test adapter concentrations and ligation time.

### *in situ* SM-Omics protocol (1)

Tissue-stained ST slides we provided as input. The ST slide was attached into the ProPlate Multi-Array slide system (GraceBioLabs, USA), with up to four ST slides fitted. The ProPlate Multi-Array system was then fixed in position by Aluminum Heat Transfer Plate (VP 741I6-GS-4, V&P Scientific, Inc, USA) on the Agilent Bravo deck. The protocol started with tissue pre-permeabilization (30 min at 33°C for mouse brain) with addition of 120μl reagent per well of exonuclease I buffer (NEB, USA). In case spleen sections were processed, the pre-permeabilization step was skipped. For complete removal of the reagents and wash solutions from the subarrays all of the robotic dispensing and aspiration steps took place in all four corners of the square wells. Pre-permeabilization reagent removal was followed by a 180μl wash in 0.1X Saline Sodium Citrate (SSC, Sigma-Aldrich, USA) at 33°C. Next, tissue permeabilization was done using 75μl 0.1% pepsin (pH 1, Sigma-Aldrich, USA) at 33°C for 10min (mouse brain) and 60min 75μl 0.1% pepsin (spleen) prepared at pH 2.5 in Tris-HCl (Sigma-Aldrich, USA). After a 180μl 0.1X SSC wash at 33°C, *in situ* cDNA synthesis reaction was performed by the addition of 75μl RT reagents: 50ng/μl actinomycin D (Sigma-Aldrich, USA), 0.5mM dNTPs (Thermo Fisher Scientific, USA), 0.20μg/μl BSA, 1 U/μl USER enzyme (both from NEB, USA), 6% v/v lymphoprep (STEMCELL Technologies, Canada), 1M betaine (#B0300-1VL, Sigma-Aldrich, USA), 1X First strand buffer, 5mM DTT, 2U/μl RNaseOUT, 20U/μl Superscript III (all from Thermo Fisher Scientific, USA). The reactions were sealed with 70μl of white mineral oil Drakerol#7 (Penreco, USA). Incubation at 30°C was performed for a minimum of 6h, after which 70μl of the released material was collected in a new 96-well PCR plate (Eppendorf, Germany). When a Cy3 fluorescent cDNA activity print was needed for tissue optimization, the 75μl *in situ* cDNA reaction mix was as follows: 50ng/μl actinomycin D (Sigma-Aldrich, USA), 0.20μg/μl BSA (NEB, USA), 1X M-MuLV buffer, 5mM DTT, 2U/μl RNaseOUT, 20U/μl M-MuLV (all from Thermo Fisher Scientific, USA), 4μl dNTP mix (dATP; dGTP and dTTP at 10mM and dCTP at 2.5mM) and 2.2μl Cy3-dCTPs (0.2mM, Perkin Elmer, USA).

### *in situ* manual ST protocol

The manual ST *in situ* protocol was performed as described in Salmén *et al[45]*. The protocol is, if not mentioned below, identical to the robotic protocol except as further described. Tissue-stained ST slide was attached in an ArrayIT hybridization chamber (ArrayIT, CA). All incubations took place on an Eppendorf Thermocycler R (Eppendorf, Germany), and reactions were covered with Microseal ‘B’ PCR Plate Seals (Biorad, CA) to avoid evaporation. Pre-permeabilization and washes were performed with 100μl reagent at 37°C and the *in situ* cDNA synthesis reaction was run without the USER enzyme, lymphoprep and betaine, at 42°C. The manual protocol then encompassed tissue removal and probe release as described[45]. Tissue removal took place in two separate steps with RLT buffer with β-mercaptoethanol and Proteinase K. 80μl of 1% β-mercaptoethanol (Sigma-Aldrich, USA) in RLT buffer (Qiagen, Germany) were added to the wells and incubated at 56°C for 1h. Following removal of the reaction mix and wash with 0.1X SSC solution, 80μl of second tissue removal mixture; 2.5μg/μl Proteinase K in PDK buffer (Qiagen, Germany) were added and the reaction was performed at 56°C for 1h. The complete reaction mix was again removed and a slide wash with one 10 minute wash of the wells with 2X SSC/0.1% SDS (Sigma-Aldrich, USA), followed by 1 minute wash with 0.2X SSC and finally 0.1X SSC was performed. Cleavage of probes from the surface was performed in the next steps and not during *in situ* cDNA synthesis. The reaction mix consisted of 1.1X Second strand buffer (ThermoFisher Scientific, USA), 0.1mM dNTPs and 1 U/μl USER enzyme (NEB, USA). 75μl of the mix was added and incubated for 3h at 37°C. The released material was collected in a new 96-well PCR plate (Eppendorf, Germany) by aspirating 70μl of the released material.

### SM-Omics library preparation (2)

Upon initiating the Agilent Bravo form the user was prompted to select either: 1, 2, 3, 4, 6 or 12 columns of the 96-well plate to run. Two positions on the Bravo deck had Peltier thermal stations (4-95°C) in the standard 96-well format. A reagent plate was prepared for robotic aspiration, transfer and dispensing of reagents. First, single-stranded cDNA was made to double-stranded material using 5 μl of the reaction mix (2.7X First strand buffer, 3.7 U/μl DNA polymerase I and 0.2 U/μl Ribonuclease H (all from ThermoFisher Scientific, USA)) for 2h at 16°C. Thereafter, the material was blunted by the addition of 5μl of 3U/μl T4 DNA polymerase (NEB, USA) for 20 minutes at 16°C. The reaction was stopped by addition of Invitrogen UltraPure 0.5M EDTA (pH 8.0, ThermoFisher Scientific, USA) to a final concentration of 20mM. The material was then purified using Ampure XP (Beckman Coulter, USA) at a bead to cDNA ratio of 1:1. Next, 27.8μl of the T7 reaction mix (46.2mM rNTPs, 1.5X T7 reaction buffer, 1.54 U/μl SUPERaseIN inhibitor and 2.3U/μl T7 enzyme; all from ThermoFisher Scientific, USA) was added and sealed with 40μl of Vapor-Lock oil (Qiagen, Germany) for an overnight 14h incubation at 37°C. After incubation, 2.1μl of nuclease-free water (ThermoFisher Scientific) was added and the Vapor-Lock was removed, followed by a bead cleanup with RNAclean Ampure XP beads (Beckman Coulter, USA) at a ratio of 1.8:1 of beads:aRNA. The material was then assessed with a Bioanalyzer RNA 6000 Pico kit (Agilent Technologies, USA). 8μl of the eluted 12μl aRNA was transferred into a new 96-well PCR plate (Eppendorf, Germany).

### SM-Omics library preparation (3)

2.5μl of either 3μM (standard) or 15μM aRNA adapters (efficient) [rApp]AGATCGGAAGAGCACACGTCTGAACTCCAGTCAC[ddC] were added to 8μl of aRNA. The reaction was then incubated at 70°C in a PCR machine for 2min and immediately chilled on wet ice. The user then again selected the number of columns they wished to run. 4.5μl T4 RNA ligation mix (3.3X T4 RNA ligase buffer, 66U/μl truncated T4 ligase 2 and 13U/μl murine RNAse inhibitor (all from NEB, USA)) were added to the aRNA/adapter solution. The ligation reaction took place at 25°C for 1h (standard) or 3h (efficient). For the SM-Omics protocol, the ligation reaction was performed for 3h in the presence of 15μM aRNA adapters. The ligation was followed by an Ampure XP (Beckam Coulter, USA) bead purification at a ratio of 1.8:1 bead:cDNA. Elution volume was 12μl. After bead purification, 2μl of a primer and dNTP mix (1:1 v/v of either 20μM or 40μM GTGACTGGAGTTCAGACGTGTGCTCTTCCGA and 10mM dNTPs) were added to the ligated samples. For the SM-Omics protocol, 40μM primer amount was added using the same volumes. Then, the samples were sealed with 40μl Vapor-Lock (Qiagen, Germany) and heated to 65°C for 5min. The Vapor-Lock was thereafter removed and 8μl of reverse transcription mix were added (2.5X First strand buffer, 13mM DTT, 5 U/μl RNaseOUT and 25 U/μl Superscript III; all from Thermo Fisher Scientific, USA), with the addition of 40μl Vapor-Lock to reseal the reaction. The samples were incubated at 50°C for 1h. 10μl of nuclease-free water was added followed by a final Ampure XP bead purification at 1.7:1 bead:cDNA ratio with a final elution of 10μl nuclease-free water.

### Staining tissues with oligonucleotide-conjugated antibodies

As described above, the fresh frozen tissue was placed on the spatial array slide and fixed at RT, followed by antibody incubations at 4°C. First, tissues were blocked and permeabilized as described above. This was followed by a series of 3 washes in 1X PBS and a last wash that was incubated for 2min. After discarding the wash, oligonucleotide-conjugated antibodies and fluorescently labeled antibodies (Biolegend, USA) were both added at a 1:100 dilution in the same buffer as in the initial permeabilization step and incubated for 1h. The tissue was then washed and the antibody conjugates fixed to the array surface in 4% PFA (Sigma-Aldrich, USA). Tissues were then fluorescently imaged and SM-Omics libraries created. The following steps were added in the library preparations to ensure collection of spatially-barcoded antibody tags. First, cDNA synthesis was performed *in situ* under the same conditions as described above. Next, second strand synthesis was also performed as described followed by an Ampure XP bead clean up as according to manufacturer’s instructions. During this clean up, material that would otherwise have been discarded after binding to the beads in standard SM-Omics library preparations, was saved and represented a population of spatially barcoded antibody tags. This elute contained short products that required a bead clean up procedure as well, where a 1.4X bead-to-material ratio was used and the final product eluted in 50μL EB (Qiagen, Germany). This material was then indexed for Illumina sequencing using Small RNA Illumina indexes in a KAPA indexing reaction as described in *Quantification, indexing and sequencing*.

### Manual ST library preparation

Manual library preparation was performed as described in Salmén *et al[45]* and included the same experimental steps as the robotic library preparation protocol, but performed manually, incubations took place in a PCR System Eppendorf Mastercycler (Eppendorf, Germany) and instead of Vapor-Lock, reactions were sealed using MicroAmp Optical 8-Cap Strips (ThermoFisher Scientific, USA). The manual procedure also included the following deviations from the robotic library preparation: T7 reaction mix of 18.6μl was used and 1.4μl of nuclease-free water was added after the 14 hours incubation.

### Manual Visium preparation

Cortical tissues from an adult mouse brain were cryosectioned at 10μm thickness and placed on Visium capture areas. The protocol was followed as in the Visium Spatial Gene Expression User Guide CG000239 Rev B as provided by 10X Genomics.

### Quantification, indexing and sequencing

qPCR library quantification and indexing were performed as described in Salmén *et al[45]*. The indexed SM-Omics cDNA libraries were diluted with 40μl of nuclease-free water to allow for a final library bead cleanup with 0.8:1 ratio Ampure XP beads to PCR products, according to the manufacturer’s protocol. Final elution was done in 16μl EB (Qiagen, Germany). Individual libraries’ fragment lengths and concentrations were evaluated on a Bioanalyzer HS (Agilent Technologies, USA) or DNA1000 Tapestation (Agilent Technologies, USA) and DNA HS Qubit assays (ThermoFisher Scientific, USA), respectively. Samples were then diluted to the desired concentration for sequencing (~1.08 pM final for NextSeq sequencing with 10% PhiX) and sequenced 27-30nt in the forward read and 55-58nt in the reverse read. For antibody tags, the final clean-up was performed at 0.9:1 ratio of beads to PCR products and elution again done in 16μl EB (Qiagen, Germany). Samples were diluted to 8pM final concentration before sequencing on an Illumina Miseq (2×25nt).

### Raw reads processing and mapping

ST, SM-Omics, Visium or antibody tag fastq reads were generated with bcl2fastq2. ST Pipeline[52] v.1.3.1 was used to demultiplex the spatial barcodes and collapse duplicate UMI sequences for ST, SM-Omics and Visium. In short, 5nt trimmed R2 was used for mapping to the mouse genome (mm10) using STAR[53]. After that, mapped reads were annotated using HTseq-count[54]. To collapse UMIs, the annotated reads needed to first be connected to a spatial barcode using a TagGD[55] demultiplexer (k-mer 6, mismatches 2). Then, UMIs mapping to the same transcript and spatial barcode were collapsed using naive clustering with one mismatch allowed in the mapping process. The output file is a genes-by-barcode matrix that was used in all further processing steps. To map antibody tags to their respective spatial barcodes, we used the tag quantification pipeline originally developed for CITE-Seq (v.1.3.2) available at https://github.com/Hoohm/CITE-seq-Count. The pipeline was run with default parameters (maximum Hamming distance of 1). We additionally provided the spatial barcodes and corrected the spatial mapping (1 mismatch) for a total of 1007 different barcodes.

### Automated image processing for spatial transcriptomics

For efficient processing, HE images were scaled to approximately 500×500 pixels using the imagemagick (https://imagemagick.org/index.php) mogrify command. In order to reconstruct the positions of all ST spots, visible (*i.e.*, not covered by the tissue section) barcode (x,y) spots were registered through “blob detection” and then refined by keeping only those “blobs” (potential grid points) that were likely to be part of a regular grid. A regular grid was then fitted to the remaining potential grid points, starting an iterative process in which the 0.1% potential grid points that least fit the grid were removed in each iteration and a new grid was fitted until the target number of grid points per row (here 35) and column (here 33) were reached. Finally, those grid points that overlapped the tissue sections were identified by building a mask that represented the tissue area and registering all grid points that were present in this mask. In order to accommodate atypical tissue coloring, bubbles, and smears present as imaging artifacts, we introduced a parameter that toggles the color channels used to detect the tissue section. Finally, an intermediate report notifies the user of irregularities in the automatic alignment process and allows for visual inspection. The output .tsv file contained barcode spots (x,y) as centroid pixel coordinates of the detected grid, as well as a TRUE/FALSE value, set as TRUE if the barcode spot was detected as under the tissue section area.

### SpoTter Integration with ST Pipeline and Quality Control (QC) reporting

The following steps integrate the output from the automated image alignment steps with the output gene-by-barcode expression file as produced by the ST Pipeline v.1.3.1. The barcode (x,y) spots approximated as under the tissue section were used for subsetting the ST Pipeline gene-by-barcode file. Then, the original H&E images were downscaled and cropped using the following imagemagick commands: convert HE_image.jpg -crop width“x”height+xa+ya; where width and height represented the Euclidean lengths between (x,y) grid detected barcode spots (33,35), (1,35) and (1,35), respectively. xa and ya were described as the centroid pixel coordinates of the grid point (33,35). The cropped H&E image was then rotated as follows: mogrify -flop -flip HE_image.jpg and this image was then used as input to the QC reporting system and for the GUI annotation tool. A final quality control (QC) report was created when running SpoTteR.

### Comparison of SpoTter *vs*. ST Spot Detector *vs*. manual alignment

To be able to compare the automated image processing developed here to that of manually processed images, we acquired an additional image of the ST array area after the experiment was performed and the tissue had been removed from the array surface. Briefly, complementary and Cy3 labeled oligonucleotides (IDT, USA) were diluted in 2X SSC with 0.05% SDS to a final concentration of 1μM. 50μl of the diluted solution was added to the array surface and incubated with shaking (50rpm) for 10min at RT. This was followed by washing the slide in 4XSCC with 0.1% SDS and 0.2X SSC. The array frame and all ST barcode positions had then efficiently been labeled and acquired on the same imaging system as described. All input images in the following comparisons were the same approximate input sizes and resolution. The ST spot detector tool previously developed[46] uses the H&E and Cy3 images as input. Due to its intrinsic scaling factor and input image size requirements, initial pre-processing of both images was needed, such that images be linearly downscaled to 30% of their original size and both images individually cropped to represent the same FOVs as collected during the imaging step. However, cropping was only needed if the user did not have the possibility to automatically acquire the same FOVs using the same starting (x,y) positions. For manual alignment, we used Adobe Photoshop for initial pre-processing, same as in the previous step. Both the H&E and Cy3 acquired images were downscaled to 30% of their original size, rotated 180 degrees and aligned to the same starting (x,y) pixel coordinates. This was followed by cropping both images along the middle of the first and last row and column. The tissue boundaries were detected using the magic wand function (32px) and the selected subtracted in the Cy3 image. Spots boundaries were again detected using the same magick wand function and the background noise cleaned up using the bucket fill function (250px) in a grayscale image. This grayscale image was further used in Fiji[56] to detect the centroid coordinates of each ST barcode spot. Following Fiji processing, we translated (x,y) pixel centroid coordinates to ST barcode spot coordinates (as given during the demultiplexing step in the ST pipeline). For SpoTeR input, we only provided the original H&E imaged as acquired by the imaging system with no GUI-based preprocessing. For speed comparisons, total time needed for preprocessing steps was measured first. For manual processing, the pre-processing steps included alignment of the H&E and Cy3 images with Adobe Photoshop 2019 and creation of an ST array spots files. For ST Detector pre-processing time, we only took into consideration the time needed to open the same images in Adobe Photoshop, downscale them to 30% size and crop them the same size without any other image handling processes performed. For SpoTteR, preprocessing included the downscaling step performed with imagemagick and incorporated into the workflow. Processing steps were then performed and time was measured as described before. Total speed was considered as 1/t [s-^1^] where t represents the sum of time needed for both the pre-processing and processing steps. False positive and negative rates were calculated as percentage of spots present or absent in SpoTteR or ST Detector as compared to manually processed spot coordinates.

### Estimating lateral diffusion

Two consecutive mouse cortex fresh frozen sections were processed. One was processed manually as described earlier[45] while the other was processed using our devised robotic liquid handling setup. For these tests, we created poly(d)T arrays in-house according to manufacturer’s instructions (Codelink, Surmodics, USA) using amine-activate slides. The surface area covered with poly(d)T probes was 6×6mm. Both the H&E and gene activity Cy3 images were processed in Fiji[56]. Cell boundaries were detected (Analyze > Plot Profile) with 10% signal intensity and these were used as breakpoints to estimate Cy3 signal diffusion as lateral diffusion. Left and right cell boundaries (detected as local maxima in respective images) representing opposite sides of each cell were used in the estimate and a total of 50 cells used in each condition. A 0.1728 pixel to distance conversion ratio was used. If a diffusion distance measure was scored as negative it implied that the Cy3 signal was contained within the detected cell boundaries, and positive if outside those same boundaries.

### Estimating reproducibility of SM-Omics *in situ* reactions

Scikit-image[57] was used to process the H&E and respective fluorescent gene expression images. First, a grayscale fluorescent image was smoothed using a Gaussian filter (sigma=0.01). Then, we applied morphological reconstruction by dilating the image edges through filtering its regional maxima. This enabled us to create a background image value that could be subtracted from the original image and used in further analysis. Then, we created an elevation map with a sobel filter to mask the elevated points. This image could then be used in a tissue (*i.e.*, object) detection step using watershedding. The inverted tissue boundaries were subtracted from the detected fluorescent tissue gene expression signals and used in all further analysis. The medians of the fluorescent signals were compared using a Wilcoxon ranked sum test.

### Image annotation

To manually annotate tissue images based on their H&E features, we used a previously adapted graphical and cloud-based user interface[26]. We assigned each ST (x,y) coordinate with one or more regional tags. The region names used to annotate MOB were: Granular Cell Layer (GR), Outer Plexiform Layer (OPL), Mitral Layer (MI), Internal Plexiform Layer (IPL) and Glomerular Layer (GL) and to annotate mouse cortex were: Cerebral nuclei (CNU), Cortical subplate (CTXsp), Fiber tracts, Hippocampal formation (HIP), Hypothalamus (HY), Isocortex (ISOCTX), Midbrain (MB), Piriform area (PIR) and Thalamus (TH). For annotating spleen, we used four major areas: Red pulp, B-follicle, Marginal zone and Periarteriolar lymphoid sheaths (PALS).

### Comparisons between spatial gene expression profiles

For comparisos between the SM-Omics and ST datasets, reads were first downampled to the same saturation level (64%; chosen based on estimated saturation curve) before invoking a ST pipeline mapper, annotator and counter run to receive UMIs per spatial (x,y) barcode as described previously. Depending on sequencing depth, a gene was counted as expressed if the corresponding transcript was present in more than 10^−6^ of the sequencing depth. The total count over all spots per gene and sample were then normalized[58]. Pearsons’s correlation coefficient between the average and normalized samples as well as the Wilcoxon’s rank-sum tests was calculated using Scipy v1.2.0[59]. To compare the performance of Visium and SM-Omics, we sequenced both libraries to an average depth of ~65 million paired end reads. For Visium, we sequenced 29 nt in the forward and 43 nt in the reverse read. Reads were downsampled to the same saturation level. Both datasets were processed using the ST pipeline as described above. Briefly, we mapped reads to the modified transcriptome reference as suggested and following instructions by 10X Genomics. Conventional GTF files used in the annotation step with HTseq-count were converted so that all transcript features now carried an exon tag used in counting transcripts. UMI collapsing was done using a naive approach and allowing for 1 low quality base present in either of the datasets. Unique molecular identifiers (UMI) per measurement were calculated as previously described[52].

### Saturation curve generation

Number of unique molecules was calculated by subsampling the same proportion of mapped and annotated reads from each sample and then ran the samples through ST Pipeline v.1.3.1, where unique molecules were calculated as previously described.

### Calculating quantitative immunofluorescence profiles per SM-Omics spot

First, we trained a random forest classifier using the Ilastik[60] framework to extract probabilities of the positive class assignment ie. positive antibody signals from our IF mouse brain images. Separate classifiers were trained to each antibody used and a total of ~10 images with at least 10 fields of view were used in the training process. In each classifier, we used two labels for classification: signal and background. Respective full-sized fluorescent microscopy images were then processed and output probabilities used in the following steps. For spleen data, raw fluorescent images were used as input in the following steps. First, images were processed as described in *Estimating reproducibility of SM-Omics in situ reactions*. Calculated background was removed from each image, signal boundaries estimated using watershedding followed by creating a binary mask image. This mask was then overlaid with the original fluorescent image and this image was then used in all following steps. To quantify the fluorescent signal intensities per ST spot, the image was cropped into a 33×35 matrix creating smaller patches; each patch sized at ±1% image from the centroid of each ST spot. Finally, the intensity from each spot area was calculated as the sum of the fluorescent signal detected in that spot patch.

### Spatial gene and antibody-based expression analysis

Statistical analysis of the spatial gene and antibody tag expression data was performed using Splotch’ one- or two-level hierarchical model as previously described[31]. In short, the model captures spatial expression in anatomical regions while accounting for experimental parameters such as, in our case, different animals, and calculates gene or antibody expression estimates for each single gene or antibody in each annotated spatial spot. To find targets which were differentially expressed in an annotated morphological region, we computed a one-*vs*-all comparison and took those values with a positive log Bayesian factor (BF). Expression estimates from Splotch were used when calculating the correlation between gene expression and antibody tag counts. The expression and counts mean were calculated per annotated region and then scaled from 0 to 1 within each sample. The correlation between gene expression and fluorescent signal was calculated in the same way, but the fluorescent signal matrix, prepared as explained in *Calculating quantitative immunofluorescence profiles per SM-Omics spot*, was used instead of the antibody tag counts matrix.

### Comparison to Allen Brain Atlas data

To validate our findings, we downloaded ISH gene expression data from four major regions; GL, GR, MI and OPL, from the Allen Brain Atlas (ABA) (https://mouse.brain-map.org/). For comparison in cortex samples, we used the following regions from ABA: piriform-amygdalar area (PAA), postpiriform transition area (TR) in addition to CNU, STXsp, HIP, HY, ISOCTX, MB and TH. Prior to enrichment analysis, genes found in PAA, TR and PIR in ABA were merged into one region name: PIR. We filtered genes with fold change >1 and expression threshold >2.5 in ABA and compared to genes with positive fold change and log(BF) in our Splotch data and computed a one-sided Fisher’s exact test using Scipy v1.2.0[59]. FDR was estimated using the Benjamini-Hochberg[61] procedure. One of the top most differentially expressed genes in both SM-Omics and ABA was chosen from each region and its expression visualized. The visualizations were compared to the corresponding *in situ* hybridization (ISH) images, downloaded from the ABA webpage. A reference ST dataset[24] was also analyzed using Splotch with the same settings as used for SM-Omics, visualized and compared to SM-Omics.

## Data and code availability

Raw and processed data will be available at the Single Cell Portal (https://portals.broadinstitute.org/single_cell/study/SCP979).

## Acknowledgments

We thank Ania Hupalowska for making all graphical illustrations. We thank Tarmö Äijö for help with Splotch and Theresa Teneyck for help with Bravo protocol implementation. Work was supported by the Knut and Alice Wallenberg Foundation, the Royal Swedish Academy of Sciences and Swedish Society for Medical Research (S.V.), the Hans Werthén Foundation (B.L), HFSP long term fellowship (LT000452/2019-L) (J.K)., the Klarman Cell Observatory, the Manton Foundation, and HHMI (A.R.). S.V is supported as a Wallenberg Fellow at the Broad Institute of MIT and Harvard.

## Author contributions

S.V. and A.R. designed the study and experiments; S.V. and B.L. performed the experiments with help from Å.S. for the automation testing steps; S.V. and J.K. designed and implemented the automatic alignment and reporting tool; B.L. analyzed data with supervision from S.V.; O.R.R. helped plan experiments. S.V., B.L. and A.R. wrote the manuscript with input from all the authors. All authors discussed the results.

## Supporting information

**FigS1.**
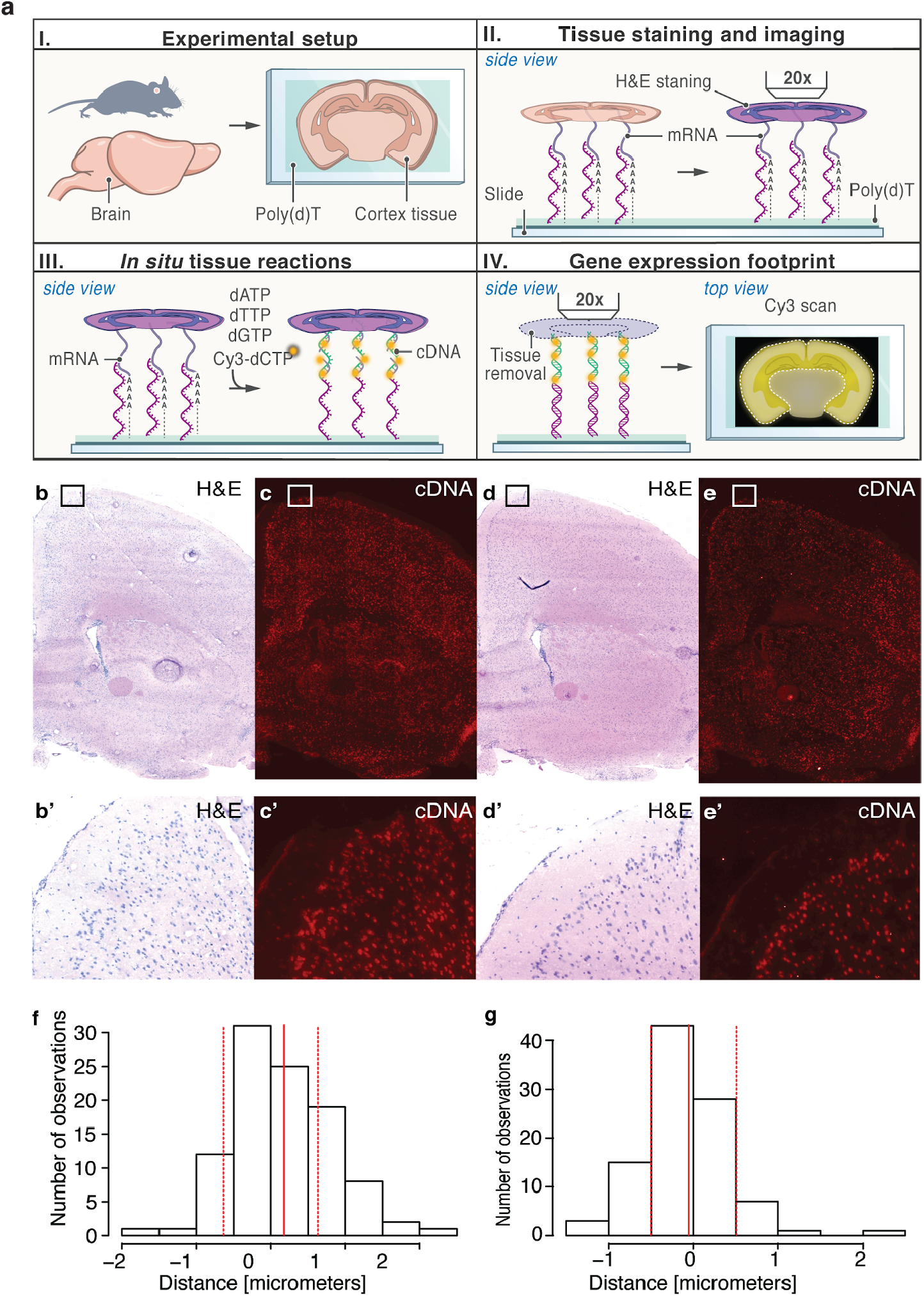
Feasibility of SM-Omics *in situ* reactions. **(a)** SM-Omics approach combines automated imaging of H&E (or IF) stained tissue sections to create spatially resolved cDNA expression footprints. First, brain sections are deposited on a mock array with poly(d)T capture area (I) and stained for H&E histology (II). Then, mRNAs are captured on the mock slide and cDNA molecules *in situ* fluorescently labeled (III) to create a spatial cDNA gene expression footprint (IV). **(b)**and **(b’)** H&E images of the cortex region on the adult mouse brain for manually prepared samples; coronal brain half and zoomed in region respectively. **(c)** and **(c’)** Fluorescent gene activity cDNA footprints corresponding to **(b)** and **(b’)**. **(d)** and **(d’)** H&E image of the adjacent cortex region processed with SM-Omics; coronal brain half and zoomed in region respectively. **(e)** and **(e’)** Fluorescent gene activity footprints corresponding to **(c)** and **(c’)**. **(f-g)** Histograms of distances between detected H&E cell boundaries and fluorescent prints for ST and SM-Omics preparations marking lateral diffusion metrics. Solid red lines represent mean and dashed lines standard deviations of the distributions (n=100).

**FigS2.**
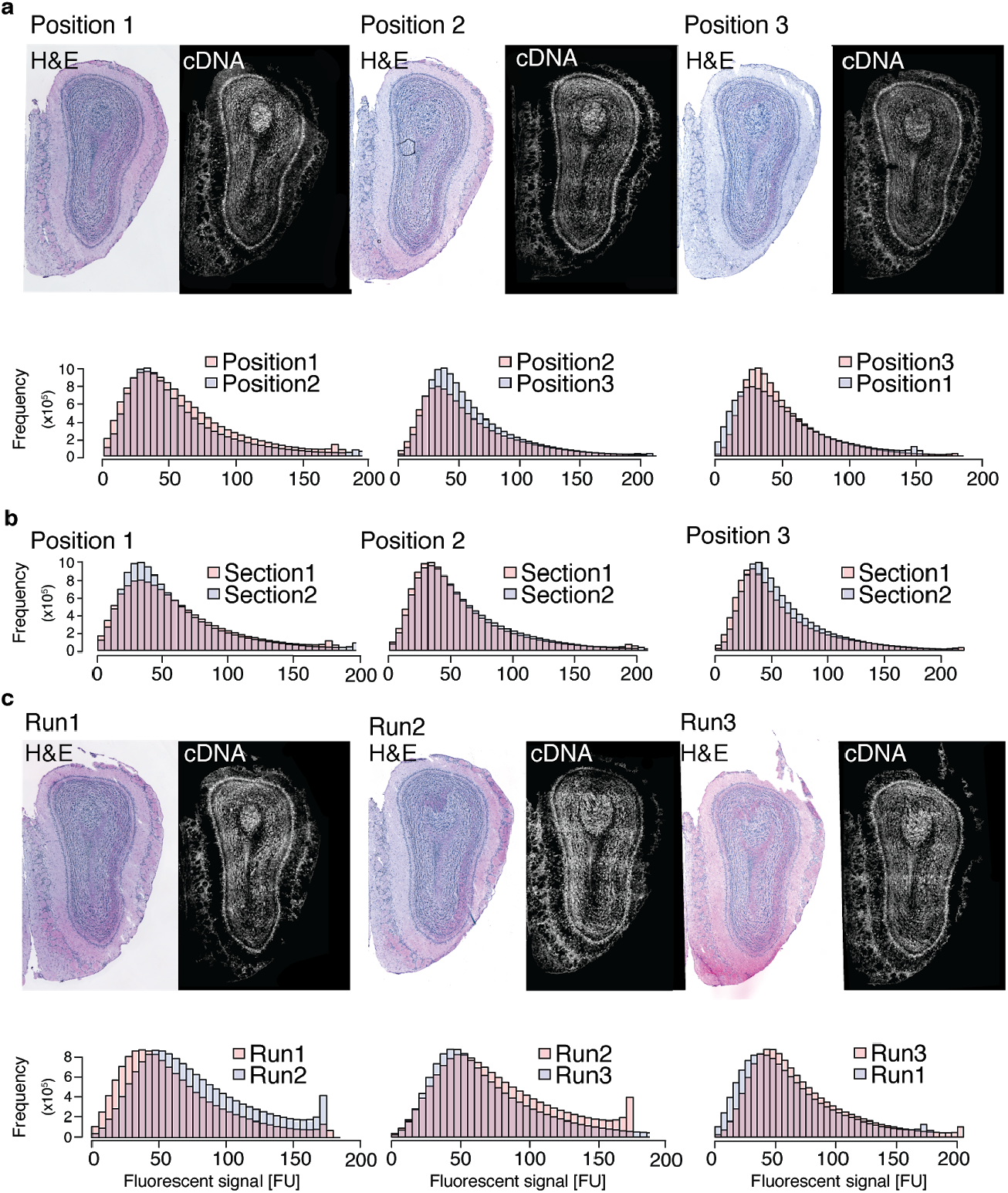
Evaluation of automated *in situ* reactions within and between SM-Omics runs on MOB tissues. **(a)**H&E images followed by detected fluorescent (cDNA) footprints (**Methods**) reflecting gene activity in the tissue sample. Each image combination (H&E and cDNA) denotes a respective position (1-3) used during one SM-Omics *in situ* optimization run (upper panel). Histograms of fluorescent tissue footprints detected in one SM-Omics run using three slide positions (lower panel). No significant differences were detected between the medians of the distributions (Wilcoxon’s rank-sum test, p>0.05). **(b)** Histograms of replicate fluorescent tissue footprints (cDNA) detected (**Methods**) in one SM-Omics run and slide position. No significant differences were detected between the medians of the distributions (Wilcoxon’s rank-sum test, p>0.05). **(c)** H&E images followed by detected fluorescent (cDNA) footprints (**Methods**) reflecting gene activity in the tissue sample. Each image combination represents a result from a separate SM-Omics run. Histograms of fluorescent tissue footprints detected between three runs. No significant differences were detected between the medians of the distributions (Wilcoxon rank-sum test, p>0.05).

**FigS3.**
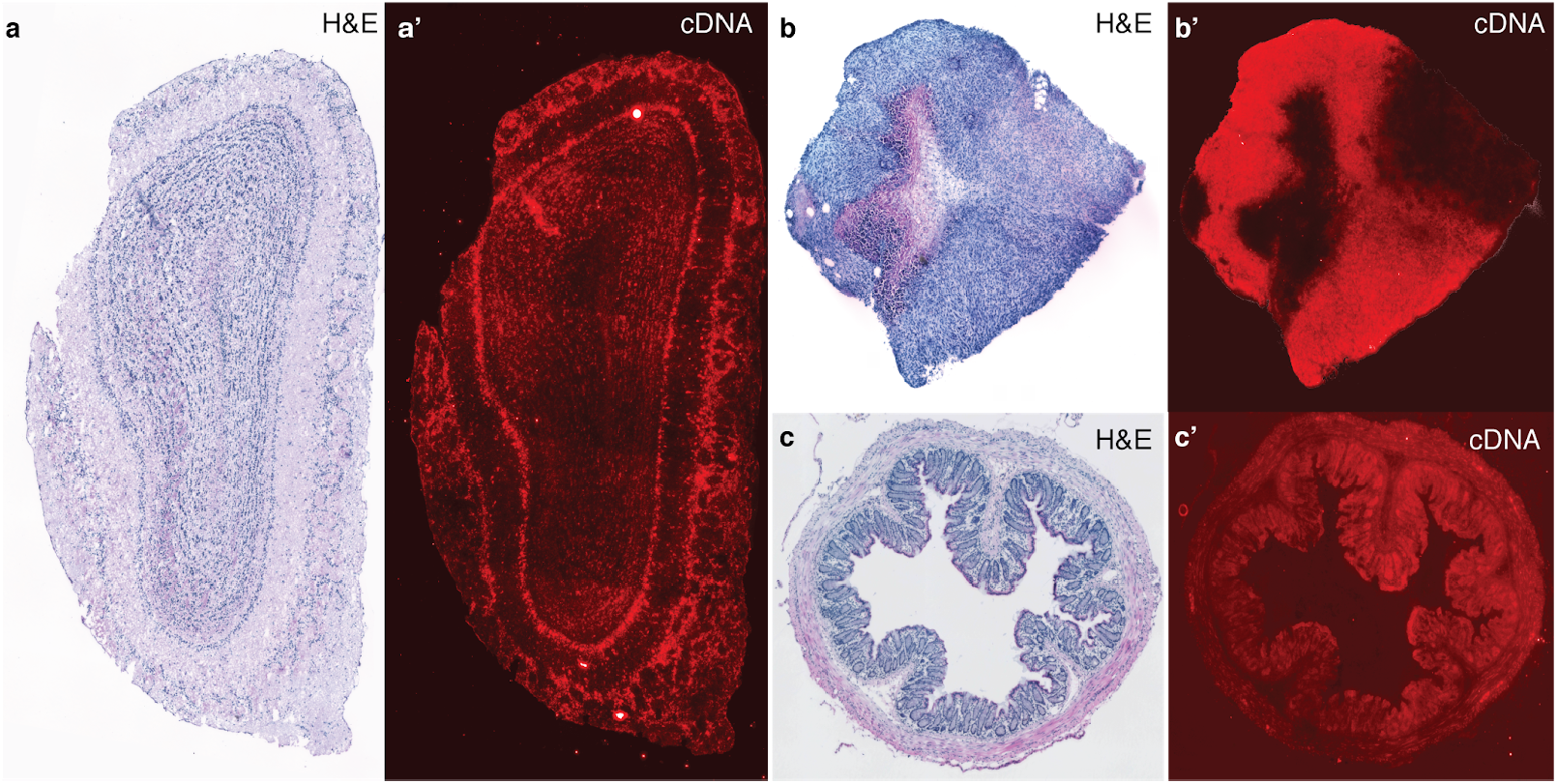
Performing SM-Omics *in situ* reactions on different tissue types. (a-a’) H&E and fluorescent print for the main olfactory bulb of the adult mouse brain. **(b-b’)** H&E and fluorescent print for the MC38-OVA inject adoed cell lines into a preclinical model of colorectal cancer. **(c-c’)** H&E and fluorescent print for the adult mouse colon.

**FigS4.**
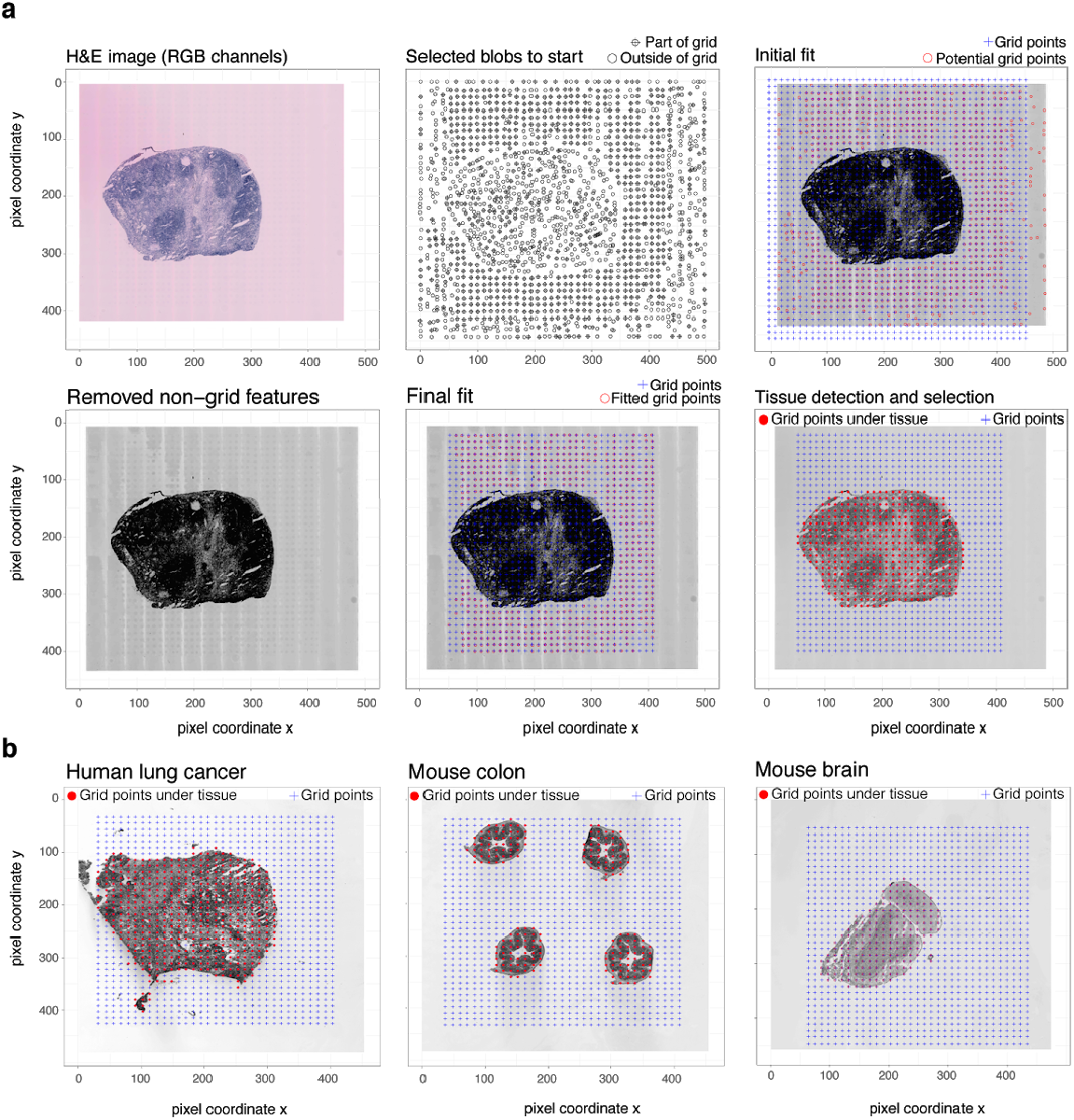
Tissue and array grid detection with SpoTteR. **(a)** The RGB tissue H&E stained image as input. The RGB image is split into 3 color channels and circular features are detected. Those features that fit a grid pattern (33×35 matrix) are used for the initial fit. Then circular features outside the grid are removed and the process of grid fitting repeated until a perfect 33×35 matrix is adjusted and positioned. Then the tissue is detected and grid spots under the tissue are easily subtracted. **(b)** SpoTteR performance for tissue and grid detection in three different tissue types: human lung cancer, mouse colon and mouse brain.

**FigS5.**
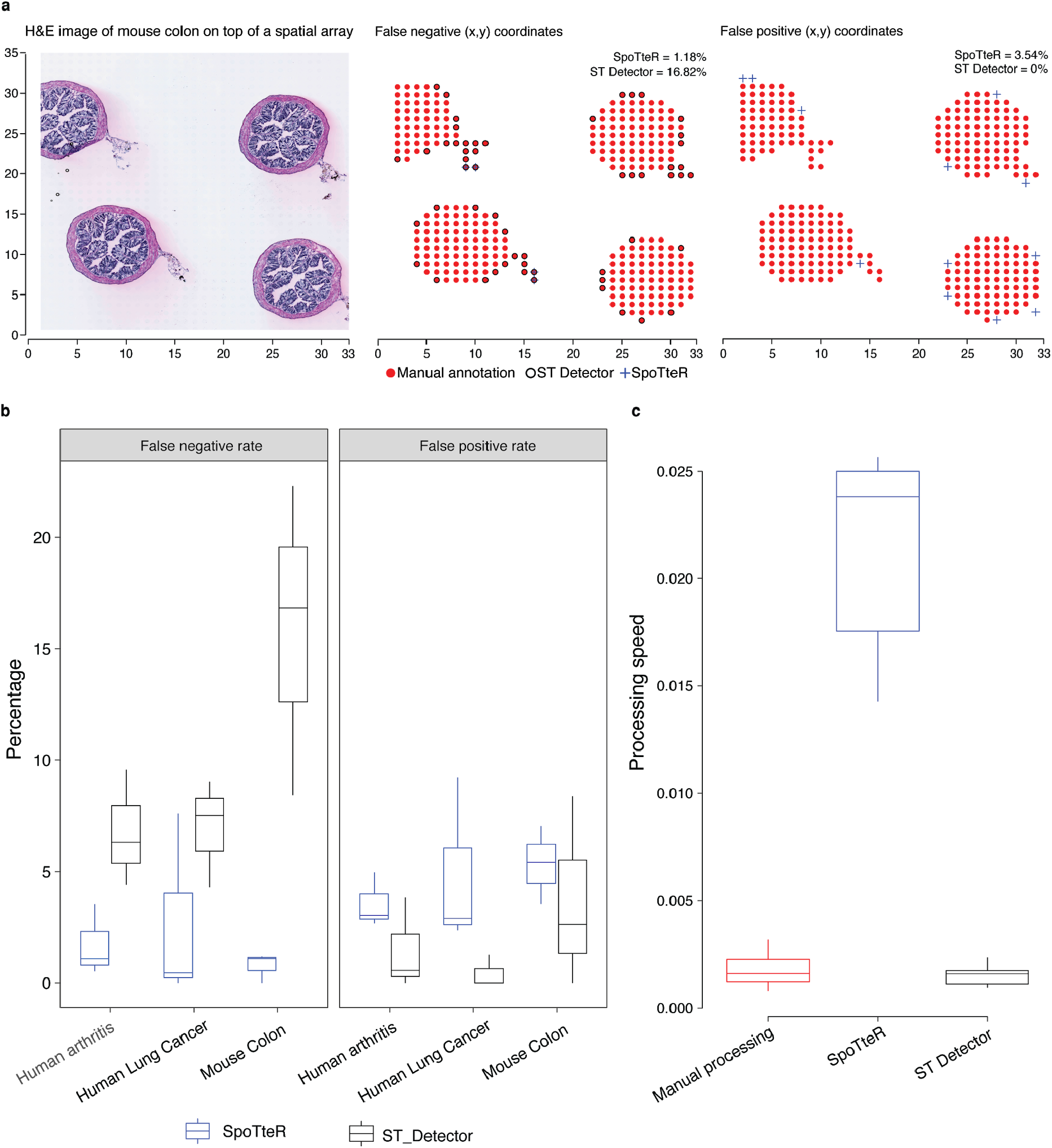
SpoTteR performance. **(a)** H&E image with corresponding false negative and positive ST barcode spot (x,y) positions using SpoTteR (blue cross) or ST Detector (black circle) as compared to the manually curated positions (filled red circle) for a mouse colon sample. **(b)** Total false negative and positive rates per processed tissue type. **(c)** Processing speed (given as 1/time [s^−1^]) for three tested processing approaches with note that there is no hands-on processing needed with SpoTteR while the other approaches require additional user input in either pre-processing or processing steps (**Methods**).

**FigS6.**
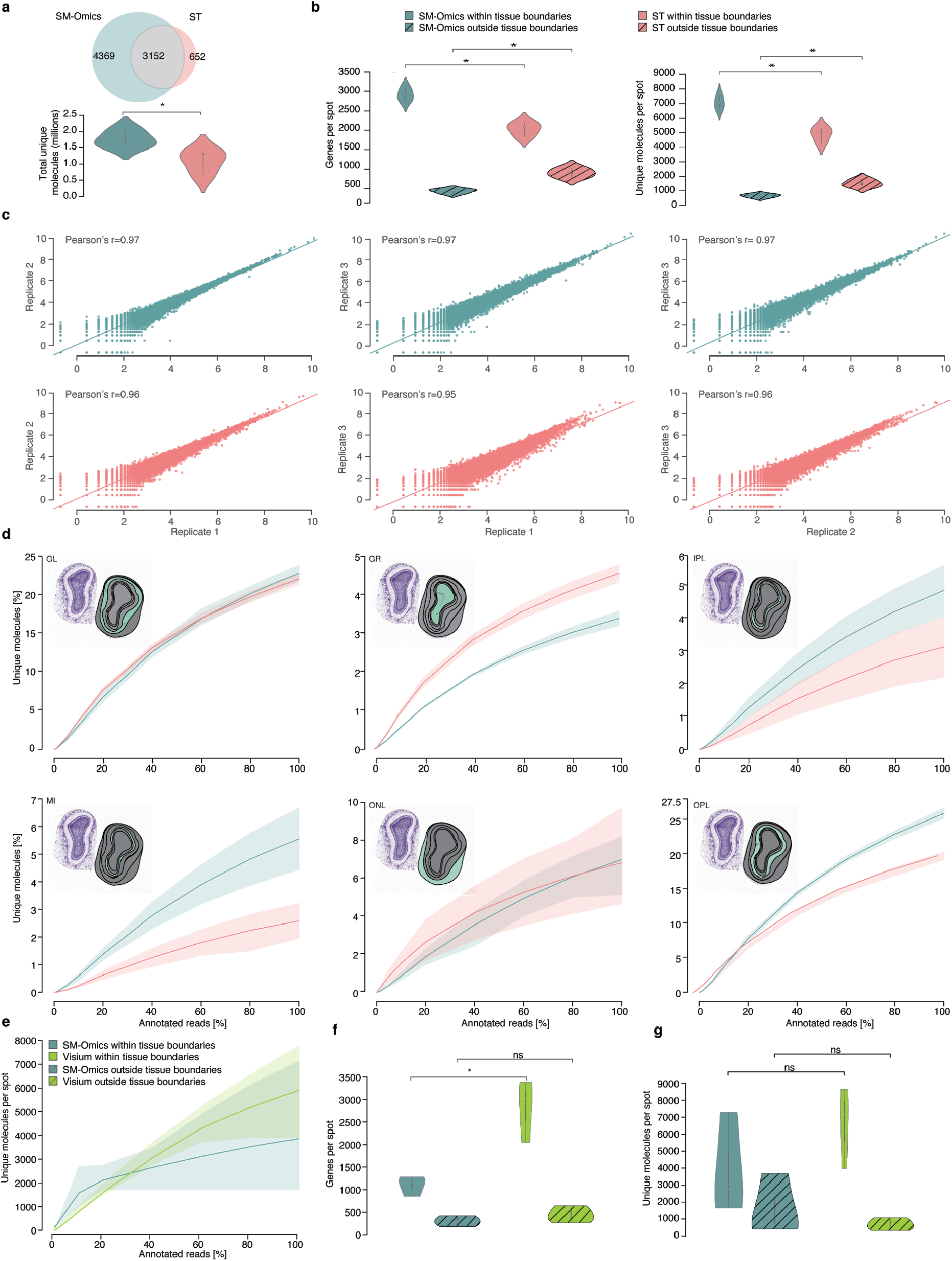
SM-Omics metrics comparisons to other array versions. **(a)** Total number of expressed genes and their intersection and total number of unique molecules detected under the tissue boundaries for SM-Omics (n=3, blue) and ST (n=3, red). **(b)** Number of expressed genes and unique molecules detected per spot under and outside of the tissue boundaries for SM-Omics (n=3, blue) and ST (n=3, red). **(c)** Correlation of the normalized pseudo-bulk gene expressions between SM-Omics (n=3) and ST (n=3). Denoted is the Pearsons’s correlation coefficient (r) between replicates. Colored line represents the linear regression line between the replicates. **(d)** Saturation curves as mean proportion value of unique molecules detected per annotated morphological region with 68% confidence interval in SM-Omics (blue line, n=3) and ST (red line, n=3). **(e)** Saturation curves (downsampled raw data, **Methods**) depicting total number of detected UMIs between SM-Omics (blue line, n=3) and Visium (green line, n=3) with 68% confidence interval. Total number of detected genes **(f)** and UMIs **(g)** per spot under and outside of the tissue boundaries in SM-Omics (n=3, blue) and Visium (green, n=3) at highest available sequencing saturation point. Annotated region abbreviations: GL (Glomerular Layer), GR (Granular Cell Layer), MI (Mitral Layer), IPL (Internal Plexiform layer) and OPL (Outer Plexiform Layer). Nissl stain and corresponding annotation regions shown in each subplot (**a-d**, positive region shown in green and rest in gray) as an example from the Allen Brain Atlas data. Color legend (a-d) is shared between the panels as denoted in (**b**). Color legend **(e-g)** is shared between the panels as denoted in **(e)**. **(a-d)** represents data from adult mouse MOB and **(e-g)** from adult mouse cortex. Statistical significance markings (Wilcoxon’s rank sum test) are displayed; `0.01<p≤0.05(*)`. Center line, median; box limits, upper and lower quartiles; whiskers, 1.5x interquartile range in **(a-b, f-g)** and density past extreme data points in **(a-b).**

**FigS7.**
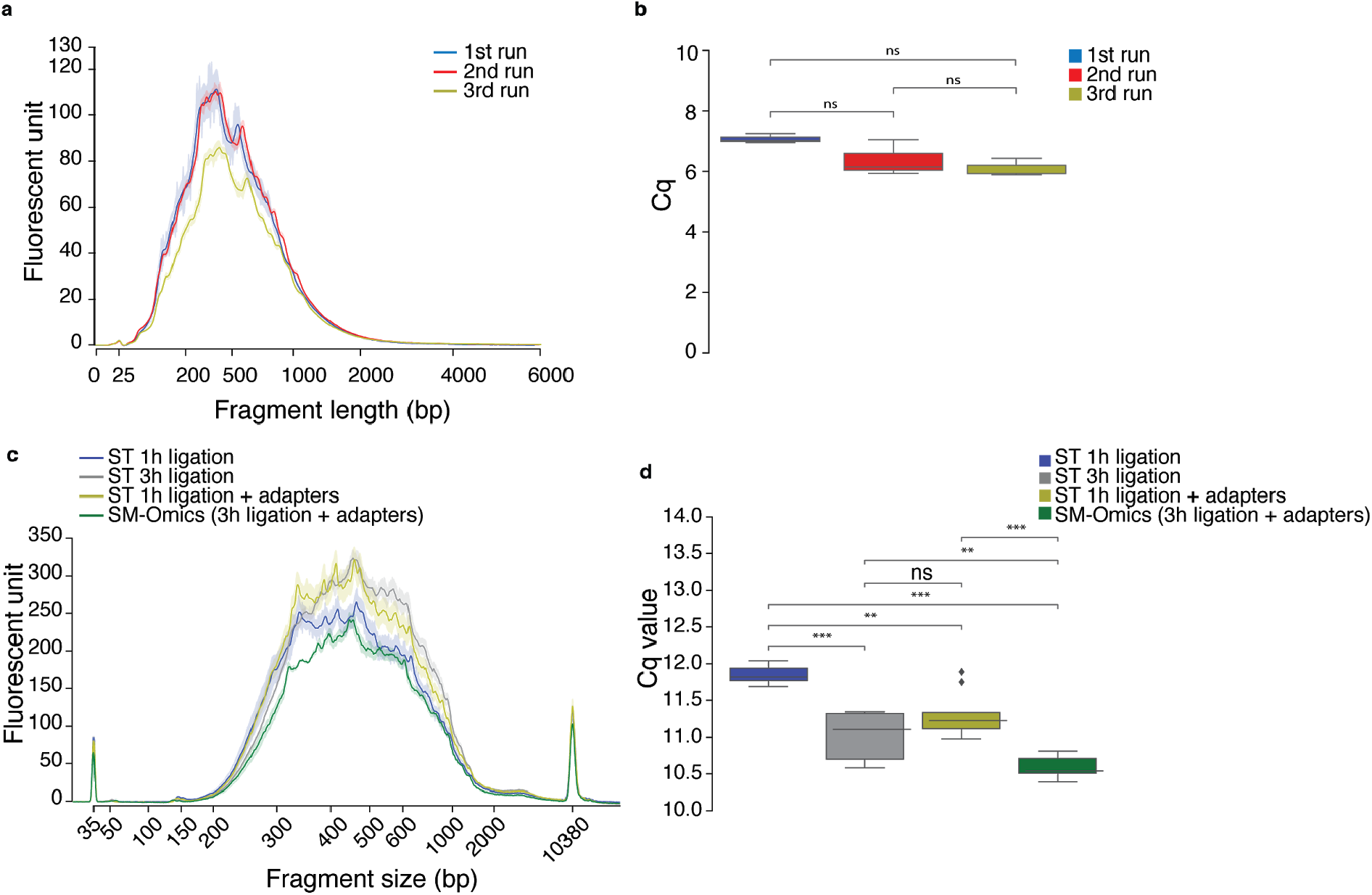
Performance of automated spatial library preparation reactions. **(a)** Mean fragment length distribution with 68% confidence interval of amplified RNA for SM-Omics samples (n=3) from three separate runs. This step represents QC results after the first part of the library preparation. **(b)** Quantitative concentrations (Cq) reflecting the final library and second part of the automatic preparation for samples processed in **(a)** from three separate runs. Results display no significant variance between the runs (p⪰0.05 (ns), Wilcoxon’s rank-sum test). **(c)** Impact of ligation reaction times and adaptor concentrations on mean fragment length distribution with 68% confidence interval of variations of final spatial libraries prepared using the automated preparation platform (n=3 for ‘‘ST 1h ligation”, “ST 3h ligation”, “SM-Omics” and n=2 for “ST 1h ligation + adapters”) **(d)** Impact of ligation reaction times and adaptor concentrations on quantitative concentrations (Cq) values for automated prepared libraries (n=9). Cq values were measured at Fluorescent unit 10,000. Statistical significance (Wilcoxon’s rank-sum test) markings are displayed: 0.05<p≤1 (ns), 0.001<p`≤`0.01 (**), 0.0001<p`≤`0.001 (***). Individual reaction conditions have been detailed in **Methods**.

**FigS8.**
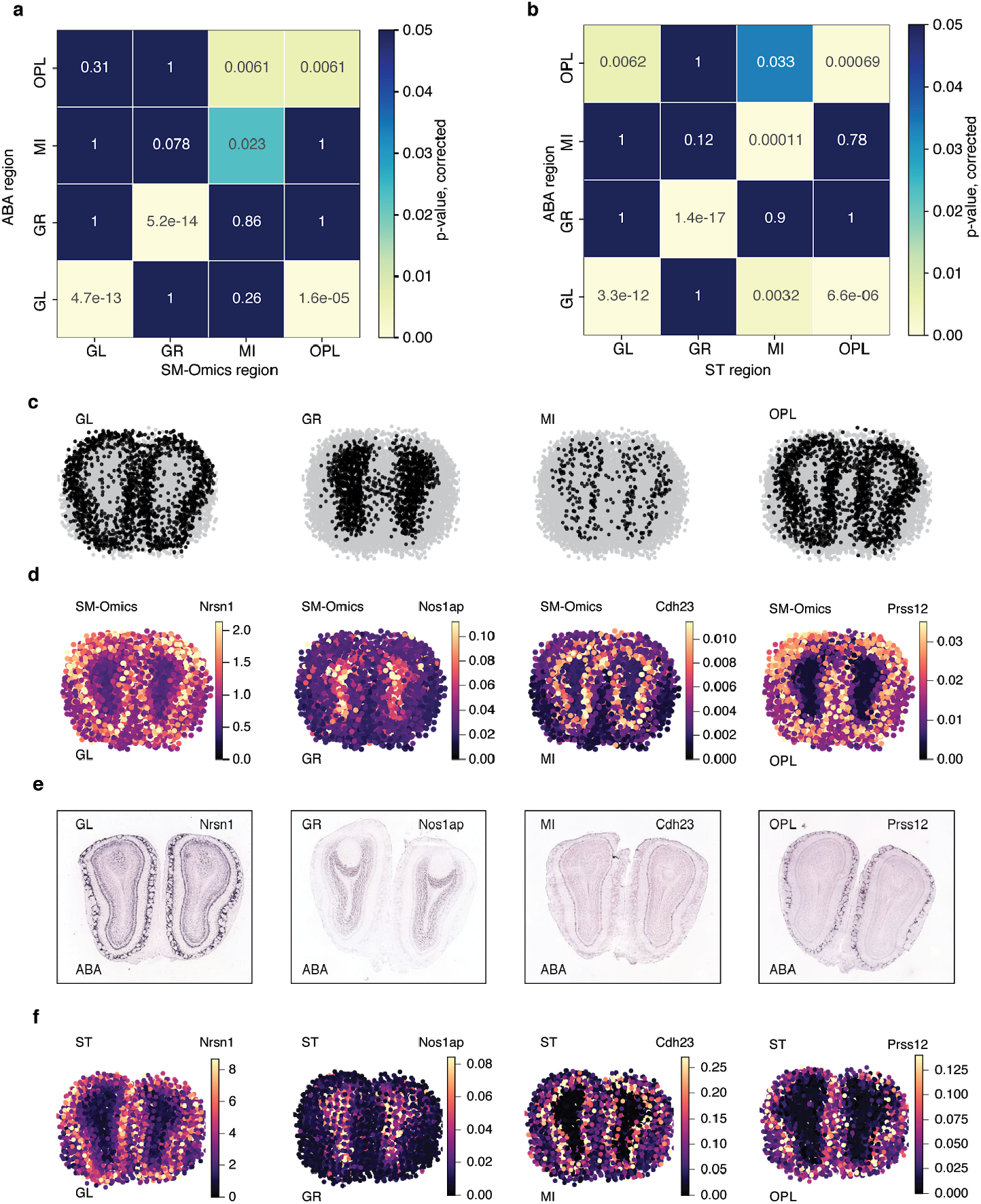
Spatial gene expression specificity and patterns in major annotated layers in SM-Omics and ST. **(a)** Morphological gene expression signatures agree between SM-Omics and the Allen Brain Atlas (ABA) for the major layers. p-value of Fisher’s test (color scale) for the enrichment of genes associated with each layer in SM-Omics (columns) and in the Allen Brain Atlas (rows). **(b)** Same as in (a) but shown for ST. Color scale denotes significant p-values (p≤0.05, Fisher’s exact test, one sided, Benjamini-Hochberg corrected for multiple testing) in panels **(a)** and **(b)**. **(c)** Examples of spatial annotation patterns (black) for four major morphological regions present in the adult MOB (**Methods**). **(d)** Examples of SM-Omics spatial gene expression patterns (color scale) for DE genes detected (**Methods**) between the regions GL, GR, MI and OPL with **(c)** corresponding ISH images from ABA and **(e)** ST spatial gene expression (color scale) for the same DE genes as in **(f).** Annotated region abbreviations: GL (Glomerular Layer), GR (Granular Cell Layer), MI (Mitral Layer) and OPL (Outer Plexiform Layer) are shared between the panels.

**FigS9.**
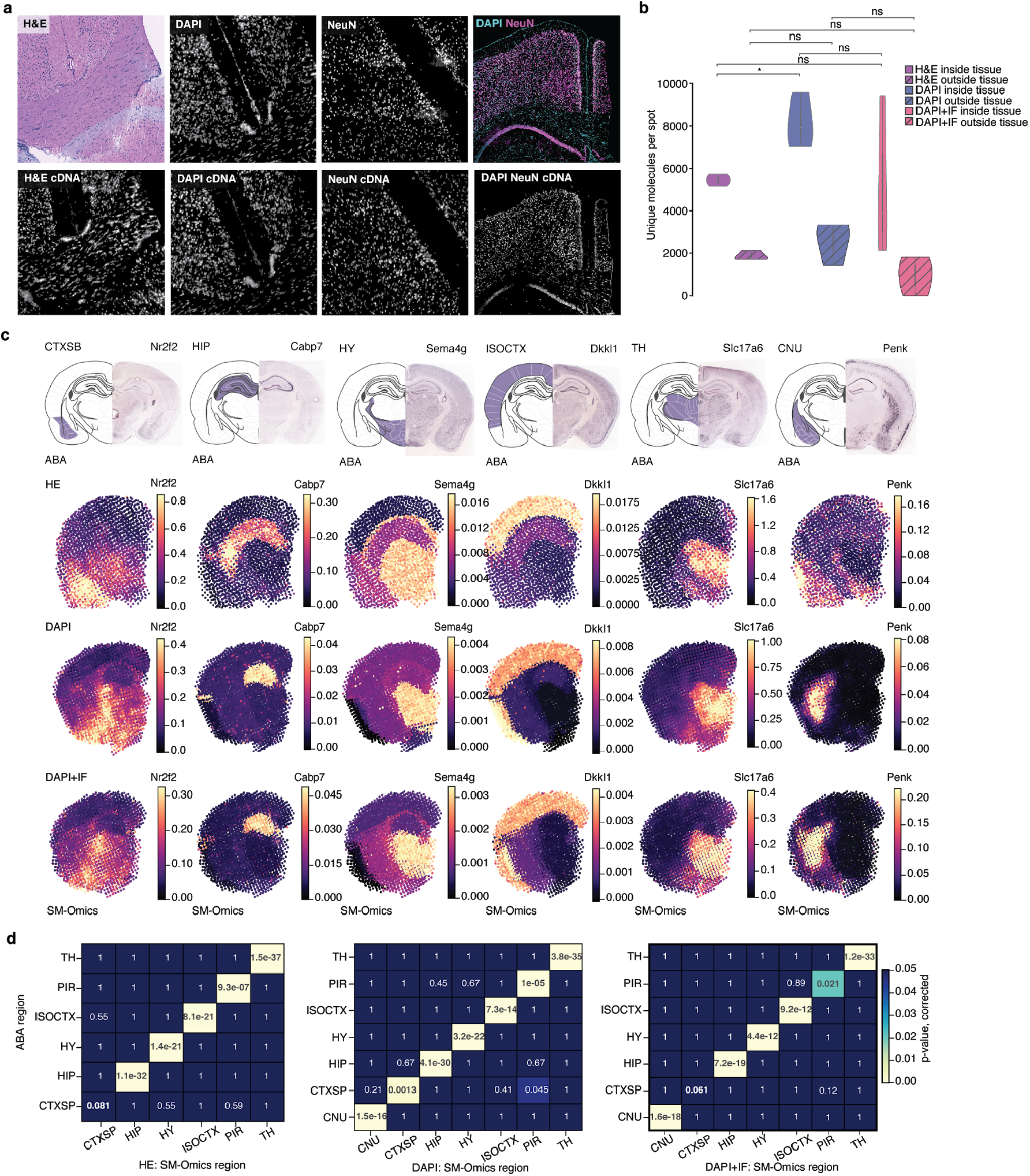
Feasibility and quality of combined antibody immunofluorescence and spatial transcriptomics measurements. **(a)** Top panel represents cortex region images in the following order: H&E stained, only DAPI stained, only NeuN stained or DAPI/NeuN stained tissues. Bottom panel shows fluorescent gene activity as Cy3 cDNA footprints corresponding to top panels. No significant differences were observed in Cy3 cDNA signal intensities between the conditions (data not shown, Wilcoxon’s rank sum test, p>0.05). Total number of detected and UMIs per spot under and outside of the tissue boundaries in SM-Omics when tissue staining was performed with 3 different conditions: H&E (purple, n=3), DAPI (blue, n=3) and a combined DAPI and immunofluorescent (IF) stain (red, n=3). **(c)** ISH images from ABA for DE genes in each morphological region (columns). Examples of SM-Omics spatial gene expression patterns (color scale) for the same DE genes detected (**Methods**). Shown in rows are spatial patterns resulting after 3 different staining conditions as in **(b). (d)** Morphological gene expression signatures agree between SM-Omics and the ABA for the all major layers present and in all 3 staining conditions. p-value of Fisher’s test (color scale) for the enrichment of genes associated with each layer in SM-Omics (staining conditions in columns) and in the Allen Brain Atlas (rows). Annotated region abbreviations: CTXSP (Cortical subplate), HIP (Hippocampal formation), HY (Hypothalamus), TH (Thalamus), CNU (Cerebral nuclei), ISOCTX (isocortex) and PIR (Piriform area) are shared between the panels.

**FigS10.**
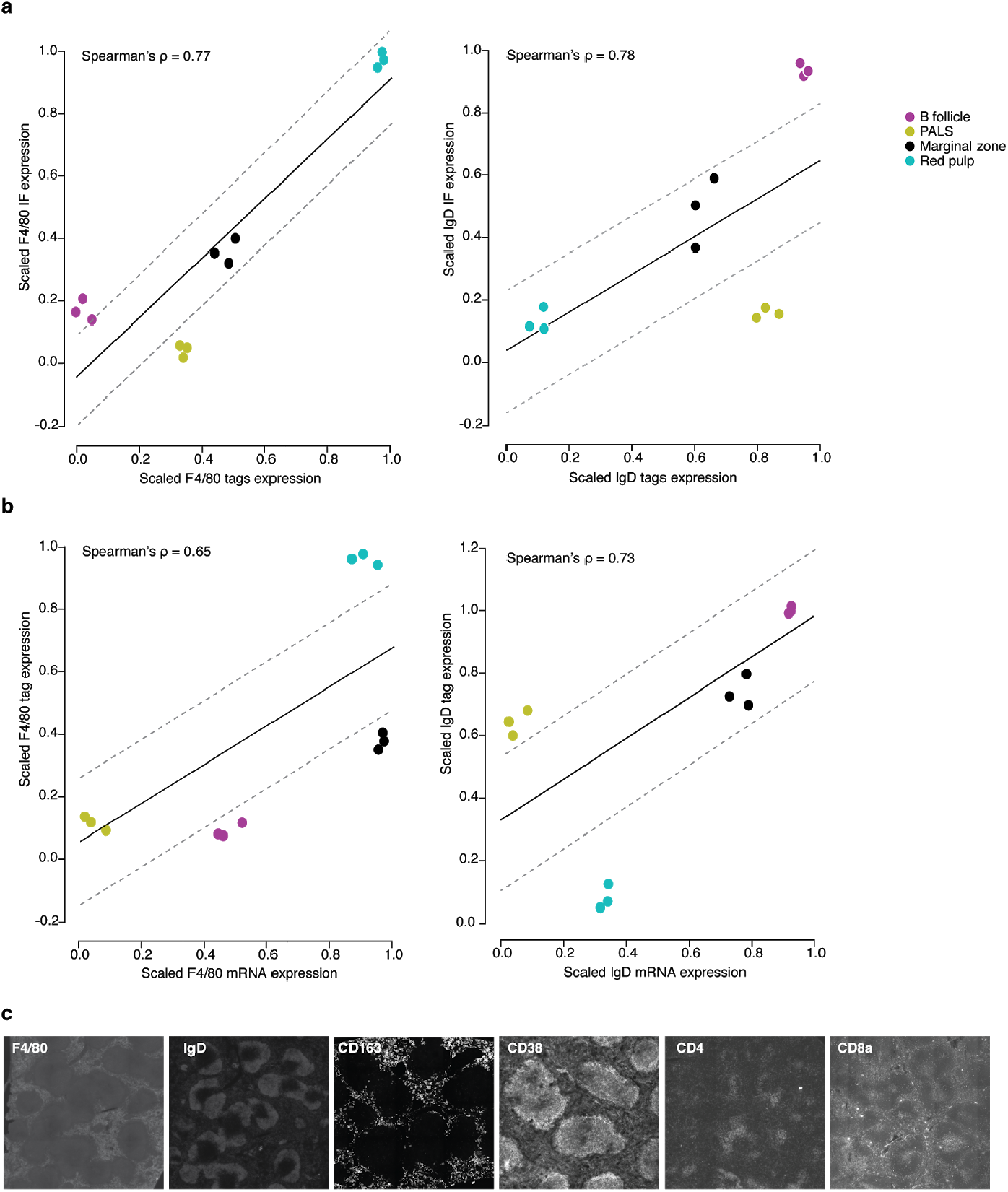
Feasibility and quality of combined antibody immunofluorescence, antibody tags and spatial transcriptomics measurements. **(a)** Correlation between scaled antibody tag and respective IF expression per tissue section ( n=3, **Methods)** for the two targets: F4/80 and IgD. Denoted is the Spearman’s correlation coefficient (ρ) between moieties. Colored line represents the linear regression line between the conditions with respective standard deviations (dashed gray lines). Color code represents 4 annotated splenic regions. **(b)** Correlation between scaled mRNA and respective antibody tag expression per tissue section ( n=3, **Methods).** Denoted is the Spearman’s correlation coefficient (ρ) between moieties. Colored line (black) represents the linear regression line between the conditions with respective standard deviations (dashed gray lines). Color code is shared with **(a). (c)** IF images of 6 antibody clones: F4/80, IgD, CD163, CD38, CD4 and CD8a.

